# The Ciliary Lumen Accommodates Passive Diffusion and Vesicle Trafficking in Cytoplasmic-Ciliary Transport

**DOI:** 10.1101/704213

**Authors:** Andrew Ruba, Wangxi Luo, Jingjie Yu, Daisuke Takao, Athanasios Evangelou, Rachel Higgins, Saovleak Khim, Kristen J. Verhey, Weidong Yang

**Author notes:** Corresponding authors: Andrew Ruba; Weidong Yang.

## Abstract

Transport of membrane and cytosolic proteins into the primary cilium is essential for its role in cellular signaling. Using single molecule microscopy, we mapped the movement of membrane and soluble proteins at the base of the primary cilium. In addition to the well-known intraflagellar transport (IFT) route, we identified two new pathways within the lumen of the primary cilium - passive diffusional and vesicle transport routes - that are adopted by proteins for cytoplasmic-cilium transport in live cells. Independent of the IFT path, approximately half of IFT motors (KIF3A) and cargo (α-tubulin) take the passive diffusion route and more than half of membrane-embedded G protein coupled receptors (SSTR3 and HTR6) use RAB8A-regulated vesicles to transport into and inside cilia. Furthermore, ciliary lumen transport is the preferred route for membrane proteins in the early stages of ciliogenesis and inhibition of SSTR3 vesicle transport completely blocks ciliogenesis. Furthermore, clathrin-mediated, signal-dependent internalization of SSTR3 also occurs through the ciliary lumen. These transport routes were also observed in *Chlamydomonas reinhardtii* flagella, suggesting their conserved roles in trafficking of ciliary proteins.

## Introduction

The primary cilium is an antenna-like projection on nearly all mammalian cells and is involved in a variety of signal transduction pathways ranging from planar cell polarity, Hedgehog signaling, and neuronal signaling to nutrient sensing, mechanosensation, olfaction, phototransduction, and cellular growth (Marshall et al., 2006, Nauli et al., 2003, Ross et al., 2005, Jones et al., 2008, Rohatgi et al., 2007, Corbit et al., Breunig et al., 2008, 2005, Gerdes and Katsanis, 2005, Scholey and Anderson, 2006, Craft et al., 2015, Wang and Dynlacht, 2018, Wheway et al., 2018, Anvarian et al., 2019, Nachury and Mick 2019). Central to all these processes is the unique composition of structural, soluble, and transmembrane (TM) proteins in primary cilia (Ostrowski et al., 2002, Pazour et al., 2005, Ishikawa et al., 2012, Breslow et al., 2018). For example, structural proteins, like tubulins and their associated proteins, form the backbone of primary cilia and facilitate mechanosensation (Singla and Reiter, 2006, Shida et al., 2010, Wloga et al., 2017), while TM proteins typically act as signal receptors (Bangs and Anderson, 2017, Hilgendorf et al., 2016, He et al., 2014, Singh et al., 2015, Barzi et al., 2011, Lerea et al., 1986, Anholt et al., 1987, Pazour et al., 2002, Yoder et al., 2002) and soluble proteins act as intracellular messengers (Nair et al., 2005, Rosenzweig et al., 2007, Jensen and Leroux, 2017, Brooks et al., 2018).

Since primary cilia are dynamic signaling organelles, the efficient cytoplasmic-ciliary transport of protein is essential for their proper function. Previous work has shown that a diffusion barrier at the base of the primary cilium restricts the entry and exit of soluble and TM proteins (Kee et al., 2012, Breslow et al., 2013, Calvert et al., 2010, Lin et al., 2013) and both active transport by microtubule-based motors and free diffusion can drive the entry of structural/soluble proteins into *Chlamydomonas reinhardtii* flagella (Hao et al., 2011, Craft et al., 2015). For TM proteins, two models have been put forth for their cytoplasm-cilium transport: 1) vesicle fusion outside the primary cilium and subsequent transport of TM proteins into primary cilia after loading onto an intraflagellar transport (IFT) train; or 2) vesicle fusion at an unknown location inside primary cilia (Figure 1A) (Jensen et al., 2004, Chuang et al., 2015, Nachury et al., 2010, Hunnicutt et al., 1990, Pazour and Bloodgood, 2008, Vieira et al., 2006). While the precise site of extraciliary fusion is unclear and may vary between cell types, the first model is largely derived from early findings in frog photoreceptors which show that rhodopsin localizes to vesicles that appear to be fusing at the periciliary ridge complex, a unique membrane domain near the base of the photoreceptors’ connecting cilium (Papermaster et al., 1985). The second model, which appears to accomplish the transport and enrichment of TM proteins in primary cilia in one step, arises from the occasional appearance of vesicle-like structures inside primary cilia and photoreceptors (Reese, 1965, Poole et al., 1985, Jensen et al., 2004, Chuang et al., 2015, Jana et al., 2018). The role of IFT in the import of TM proteins is less clear in this model.

**Figure 1.**
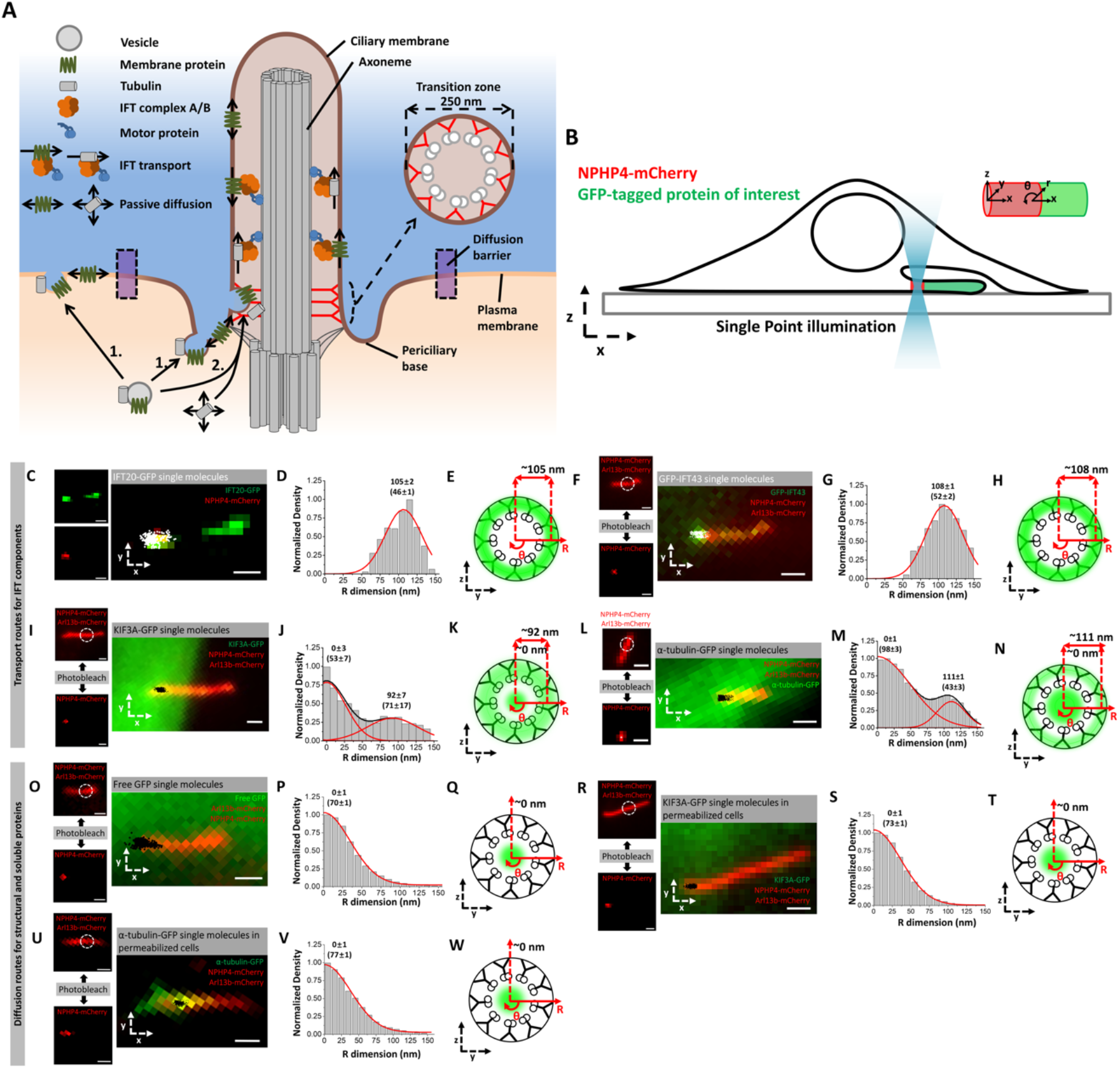
Transport routes of IFT, structural, and soluble proteins in live and permeabilized cells. **A)** Schematic highlighting the structure of TZ and two current models for the transport of structural and membrane proteins into primary cilia. **B)** Schematic of SPEED microscopy-based imaging of GFP-labeled proteins moving through the TZ labeled by NPHP4-mCherry. The spatial distributions of transiting molecules are described by both Cartesian (x,y,z) and cylindrical (x, *θ*, r) coordinate systems. **(C-E)** Imaging of IFT-B component. **C)** Epifluorescence microscopy image of IFT20-GFP (green) and NPHP4-mCherry (red) overlaid with single-molecule IFT20-GFP locations (white spots). Scale bar: 1 µm. **D)** Single-molecule IFT20-GFP locations were converted to 3D locations and plotted as a histogram along the R dimension in the cylindrical system. Numbers indicate radial distance as mean±s.e. **E)** Spatial representation of the histogram in (D) as cross-sectional view of IFT20-GFP’s transport route (green cloud) in the TZ overlaid on the ultrastructure of the TZ. **(F-H)** Imaging of IFT-A component. **F)** Epifluorescence microscopy image of GFP-IFT43 (green), NPHP4-mCherry (red), and Arl13b-mCherry (red) overlaid with single-molecule GFP-IFT43 locations (white). Dashed white circle indicates region of photobleaching to remove Arl13b-mCherry fluorescence. Scale bar: 1 µm. Small images on the left throughout this figure show the result of photobleaching Arl13b-mCherry to reveal the precise location of NPHP4-mCherry. Scale bar: 1 µm. **G)** Single-molecule GFP-IFT43 locations along the R dimension. **H)** Spatial representation of the histogram in (G). **(I-K)** Imaging of IFT motor. **I)** Epifluorescence microscopy image of KIF3A-GFP (green), NPHP4-mCherry (red), and Arl13b-mCherry (red) overlaid with single-molecule KIF3A-GFP locations (white). Scale bar: 1 µm. **J)** Single-molecule KIF3A-GFP locations along the R dimension. **K)** Spatial representation of the histogram in (J). **(L-N)** Imaging of IFT cargo. **L)** Epifluorescence microscopy image of α-tubulin-GFP (green), NPHP4-mCherry (red), and Arl13b-mCherry (red) overlaid with single-molecule α-tubulin-GFP locations (black). Scale bar: 1 µm. **M)** Single-molecule α-tubulin-GFP locations along the R dimension. **N)** Spatial representation of the histogram in (M). **(O-Q)** Imaging of GFP. **O)** Epifluorescence microscopy image of Arl13b-mCherry (red) and NPHP4-mCherry (red) overlaid with single-molecule GFP locations (black). Scale bar: 1 µm. **P)** Single-molecule GFP locations along the R dimension. **Q)** Spatial representation of the histogram in (P). **(R-T)** Imaging of IFT motor in permeabilized cells. **R)** Epifluorescence microscopy image of Arl13b-mCherry (red) and NPHP4-mCherry (red) overlaid with single-molecule KIF3A-GFP locations (black) in permeabilized cells. Scale bar: 1 µm. **S)** Single-molecule KIF3A-GFP locations along the R dimension. **T)** Spatial representation of the histogram in (S). **(U-W)** Imaging of IFT cargo in permeabilized cells. **U)** Epifluorescence microscopy image of Arl13b-mCherry (red) and NPHP4-mCherry (red) overlaid with single-molecule α-tubulin-GFP locations (black) in permeabilized cells. Scale bar: 1 µm. **V)** Single-molecule α-tubulin-GFP locations along the R dimension. **W)** Spatial representation of the histogram in (V).

From an ultrastructural perspective, primary cilia are composed of three main sub-regions. First, the basal body, which contains the 9 triplet microtubules of the mother centriole and their associated distal and subdistal appendages. Distal to the basal body is the transition zone (TZ, 300-1000 nm in length and 160-250 nm in diameter) where the 9 microtubule triplets transition to 9 microtubule doublets (termed 9+0) and is the proposed location for the components of the selectivity barrier (Yang et al., 2015, Czarnecki and Shah, 2012, Craige et al., 2010, Kee et al., 2012, Najafi et al., 2012). The TZ contains Y-shaped structures in electron micrographs that appear to tether the microtubules to the encompassing ciliary membrane. Third, the main ciliary shaft, composed of the 9+0 microtubule axoneme, is 200-250 nm in diameter and extends into the extracellular environment (Czarnecki and Shah, 2012), where it receives and transmits signals via TM receptors (Ye et al., 2018, Hilgendorf et al., 2016) and ectosomal release (Wang et al., 2014, Phua et al., 2017, Nager et al., 2017), respectively. Encompassing both the TZ and the ciliary shaft is a membrane with a unique lipid and protein composition (Yang et al., 2015, Praetorius, 2014, Molla-Herman et al., 2010, Nauli et al., 2003, Praetorius and Spring, 2003, Kaneshiro, 1987). While the primary cilium axoneme is generally modelled as a rigid 9+0 structure, various asymmetries may occur, such as a microtubule doublet turning into a singlet and/or collapsing toward the inner lumen, especially toward the distal end (Rogowski et al., 2013, O’Hagan et al., 2017, Sun et al., 2019).

To parse out the localization of TM proteins at the three main landmark regions of the primary cilium (basal body, transition zone, and ciliary shaft), we used single-point edge-excitation sub-diffraction (SPEED) microscopy, a high-speed single-molecule imaging technique developed in our lab (Ma and Yang, 2010, Ruba et al., 2018). It relies on high spatiotemporal resolution (2 ms, 10-20 nm) capture of 2D single-molecule locations of labeled proteins in live cells and the subsequent back-projection algorithm to obtain the 3D probability spatial density distribution of targeted proteins inside the primary cilium (Figure S6) (Ma and Yang, 2010, Ma et al., 2016, Ruba et al., 2017, Ruba et al., 2018). We used this approach to determine the contributions of IFT and free diffusion to the movement of soluble proteins within the lumen of the cilium shaft (Luo et al 2017). Here, we probe the transit of soluble and TM proteins at the TZ and demonstrate that the lumen is used to mediate vesicular trafficking of TM proteins as well as passive diffusion of structural and soluble proteins. We find that the axonemal lumen is the preferred route for TM proteins during two functionally-critical ciliary stages: ciliogenesis and clathrin-mediated, signal-dependent internalization. Furthermore, this luminal transport route of TM proteins is distinct from passively diffusing ciliary structural and soluble proteins. Lastly, we show that the luminal route is also present in flagella of the model system, *Chlamydomonas reinhardtii*, suggesting a wider evolutionary origin.

## Results

### SPEED microscopy maps the transit routes of ciliary proteins at the TZ

We began by validating SPEED microscopy to track protein localization in the TZ of live primary cilia by determining the transport routes for proteins whose localizations are already known. Namely, we looked at IFT components (IFT20 and IFT43), freely diffusing soluble molecules (free GFP), IFT motors and cargos (KIF3A and *α*-tubulin), and an externally-labeled TM protein (SSTR3). We chose the NIH-3T3 cell line because serum starvation causes cell cycle arrest and ciliogenesis in the majority of the cells (Ott and Lippincott-Schwartz, 2012).

Experimentally, we localized the position of the TZ in live primary cilia by detecting the fluorescence of NPHP4-mCherry, a TZ marker that localizes to the Y-shaped linkers (Awata et al., 2014), and then pre-photobleaching the fluorophore-labeled protein-of-interest by SPEED microscopy to locally reduce the labeled concentration of that protein in the TZ (Figure 1 B, Figure S1 A). Then, a high-speed CCD camera set at 500 frames per second (2 ms/frame) was used to record the 2D localizations of single, labeled proteins-of-interest in the TZ after they entered the photobleached area from neighboring regions (Video 1). We typically recorded 30,000-60,000 frames within 1-2 minutes and found the spatial shift of primary cilium is < 10 nm during the detection time. Furthermore, thousands of single molecule events with localization precisions of 10-20 nm were selected from these frames. These single-molecule localizations were then superimposed on the NPHP4-mCherry fluorescence spot used to mark the TZ to generate a 2D high-resolution spatial distribution of the protein-of-interest (Figure S1 B,C). Finally, we utilized the 2D-to-3D transformation algorithm to obtain the 3D spatial density histogram of the protein-of-interest along the radius of the primary cilium (Figure S1 D-F) (Ma and Yang, 2010, Ruba et al., 2017, Ruba et al., 2018, Ruba et al., 2018).

For the 3D reconstructed back-projection to validly represent the actual 3D distribution of single molecules, the single molecule locations must be distributed with rotational symmetry. Electron microscopy (EM) studies of the primary cilium validate the rotational symmetry of the ciliary ultrastructure, which provides a scaffolding for protein transport routes to occur (Czarnecki and Shah, 2012). Through Monte Carlo simulation, given typical experimental parameters (single molecule localization precision, number of single molecule locations, imperfect degree of labeling symmetry and rotational symmetry), we are able to obtain < 10 nm standard error on the mean peak position of routes for targeted proteins in primary cilia (Figure S7-11, Table 1). The code for the simulations, 2D-to-3D transformation, and sample and experimental data is available at https://github.com/andrewruba/YangLab.

**Table 1.** Transport route localization precision and precision/radius ratio for all 3D transport routes.

| Protein | Cilia location | Condition | # of cilia | Route mean (nm) | Route S.D. (nm) | # of single molecules | Average precision (nm) | Precision/radius ratio | $\sigma_{TR}$ (nm) |
| --- | --- | --- | --- | --- | --- | --- | --- | --- | --- |
| α-tubulin | TZ | Normal | 5 | 0 | 49 | 579 | 16 | >>0.63 | <0.1 |
|  |  |  |  | 111 | 22 | 517 | 16 | 0.14 | 3.2 |
| SSTR3 | TZ | Normal | 6 | 55 | 31 | 1471 | 18 | 0.32 | 2.0 |
|  |  |  |  | 129 | 17 | 1152 | 18 | 0.14 | 3.2 |
| IFT20 | TZ | Normal | 5 | 105 | 23 | 1162 | 16 | 0.15 | 1.8 |
| IFT43 | TZ | Normal | 5 | 108 | 26 | 1078 | 16 | 0.15 | 1.8 |
| KIF3A | TZ | Normal | 5 | 0 | 27 | 613 | 16 | >>0.63 | <0.1 |
|  |  |  |  | 92 | 36 | 633 | 16 | 0.17 | 2.7 |
| GFP | TZ | Normal | 5 | 0 | 33 | 1135 | 16 | >>0.63 | <0.1 |
| α-tubulin | TZ | Perm. | 6 | 0 | 39 | 808 | 16 | >>0.63 | <0.1 |
| KIF3A | TZ | Perm. | 5 | 0 | 37 | 918 | 16 | >>0.63 | <0.1 |
| SSTR3 | TZ | Perm. | 5 | 44 | 33 | 1137 | 23 | 0.53 | 4.2 |
|  |  |  |  | 126 | 17 | 509 | 23 | 0.18 | 4.9 |
| SSTR3 | TZ | Membrane | 4 | 131 | 12 | 1142 | 13 | 0.10 | 1.7 |
| SSTR3 | BB | Normal | 4 | 55 | 27 | 660 | 23 | 0.42 | 3.9 |
|  |  |  |  | 120 | 20 | 159 | 23 | 0.19 | 7.6 |
| SSTR3 | Shaft | Normal | 5 | 55 | 25 | 3117 | 15 | 0.27 | 1.0 |
|  |  |  |  | 131 | 22 | 2078 | 15 | 0.11 | 1.4 |
| HTR6 | BB | Normal | 4 | 45 | 26 | 765 | 27 | 0.59 | 10.3 |
|  |  |  |  | 127 | 20 | 187 | 27 | 0.21 | 9.1 |
| HTR6 | TZ | Normal | 5 | 39 | 34 | 1364 | 24 | 0.62 | 9.2 |
|  |  |  |  | 130 | 24 | 337 | 24 | 0.19 | 6.1 |
| HTR6 | Shaft | Normal | 5 | 46 | 27 | 1659 | 22 | 0.48 | 2.4 |
|  |  |  |  | 122 | 20 | 457 | 22 | 0.18 | 4.6 |
| RAB8A | TZ | Normal | 4 | 50 | 38 | 2326 | 13 | 0.26 | 0.9 |
|  |  |  |  | 133 | 20 | 402 | 13 | 0.10 | 2.9 |
| RAB8A | Shaft | Normal | 4 | 53 | 34 | 2648 | 14 | 0.26 | 1.3 |
|  |  |  |  | 134 | 16 | 669 | 14 | 0.10 | 2.8 |
| SSTR3 | TZ | 1 μM GCA | 3 | 53 | 31 | 957 | 20 | 0.37 | 2.4 |
|  |  |  |  | 130 | 22 | 372 | 20 | 0.15 | 4.4 |
| SSTR3 | TZ | 5 μM GCA | 4 | 60 | 35 | 824 | 22 | 0.37 | 4.1 |
|  |  |  |  | 136 | 17 | 424 | 22 | 0.16 | 4.6 |
| SSTR3 | TZ | 1 hr growth | 3 | 42 | 26 | 1269 | 21 | 0.50 | 3.1 |
| SSTR3 | TZ | 3 hr growth | 4 | 49 | 37 | 1596 | 14 | 0.29 | 1.6 |
|  |  |  |  | 135 | 43 | 239 | 14 | 0.10 | 3.9 |
| SSTR3 | TZ | 6 hr growth | 3 | 50 | 28 | 1122 | 21 | 0.42 | 3.2 |
|  |  |  |  | 133 | 27 | 230 | 21 | 0.16 | 5.1 |
| SSTR3 | TZ | 12 hr growth | 3 | 40 | 28 | 742 | 35 | 0.88 | 14.3 |
|  |  |  |  | 134 | 25 | 121 | 35 | 0.26 | 17.4 |
| HTR6 | TZ | 1 hr growth | 3 | 41 | 36 | 1132 | 19 | 0.45 | 2.5 |
| HTR6 | TZ | 3 hr growth | 3 | 50 | 30 | 729 | 14 | 0.28 | 2.0 |
|  |  |  |  | 127 | 21 | 182 | 14 | 0.11 | 4.1 |
| HTR6 | TZ | 6 hr growth | 4 | 35 | 32 | 876 | 19 | 0.56 | 4.7 |
|  |  |  |  | 125 | 24 | 341 | 19 | 0.15 | 4.7 |
| HTR6 | TZ | 12 hr growth | 3 | 45 | 28 | 1134 | 23 | 0.50 | 4.0 |
|  |  |  |  | 124 | 20 | 534 | 23 | 0.19 | 4.2 |
| SSTR3 | TZ | 10 μM sst | 3 | 47 | 40 | 942 | 19 | 0.40 | 3.0 |
|  |  |  |  | 137 | 19 | 143 | 19 | 0.14 | 7.9 |
| SSTR3 | TZ | 100 μM sst | 1 | 22 | 26 | 1476 | 22 | 1.0 | 8.8 |
| SSTR3 | TZ | 100 μM sst<br>+ 30 μM PS2 | 2 | 48 | 28 | 393 | 23 | 0.48 | 5.4 |
| SSTR3 | TZ | 100 μM sst<br>+ 30 μM PS2 | 2 | 131 | 24 | 138 | 23 | 0.18 | 7.8 |
| β-tubulin | Flagella | Normal | 6 | 11 | 27 | 1960 | 21 | 1.91 | 10.6 |
|  |  |  |  | 75 | 37 | 1058 | 21 | 0.28 | 2.6 |
| PKD2 | Flagella | Normal | 2 | 63 | 25 | 185 | 54 | 0.86 | 25.7 |
|  |  |  |  | 134 | 17 | 125 | 54 | 0.4 | 27.8 |
| IFT54 | Flagella | Normal | 2 | 79 | 36 | 1330 | 17 | 0.22 | 1.8 |
| KAP | Flagella | Normal | 2 | 29 | 12 | 164 | 25 | 0.86 | 10.9 |
|  |  |  |  | 77 | 28 | 147 | 25 | 0.32 | 7.2 |

Using SPEED microscopy, we first determined the 3D transport routes of components of the IFT A and B subcomplexes: IFT43 and IFT20, respectively (Follit et al., 2006, Arts et al., 2011, Hirano et al., 2017). EM work has shown that IFT occurs between the axonemal microtubules and ciliary membrane in *Chlamydomonas* flagella (Rosenbaum and Witman, 2002, Kozminski et al., 1995). Therefore, we expected the 3D density histograms of IFT43 and IFT20 to lie in this space. Indeed, the transport routes for IFT20-GFP and GFP-IFT43 were found at 105±2 and 108±1 nm along the cilium radius with route widths (defined as ±1 standard deviation about the peak position in Gaussian function) of 46±1 nm and 52±2 nm (Figure 1 C-H). Compared to the ciliary dimensions as determined by EM, these IFT components localize between the microtubules and ciliary membrane.

Next, we verified the localizations of IFT motors, IFT cargos, and freely diffusing small molecules. For the IFT motor, we determined the transport route for KIF3A-GFP, a component of the heterotrimeric kinesin-2 (KIF3A/KIF3B/KAP) complex that drives anterograde IFT (Engelke et al., 2019, Pazour and Rosenbaum, 2002, Kozminski et al., 1995). In agreement with our previous work in the cilium shaft (Luo et al., 2017), KIF3A was found to have two transport routes at the TZ: an outer route at 92±7 nm along the cilium radius with a route width of 71±17 nm, and an inner route at 0±3 nm along the cilium radius with a 52±2 nm route width (Figure 1 I-K). The IFT cargo α-tubulin-GFP (Hao et al., 2011, Craft et al., 2015) also occupies two distinct transport routes in the TZ: one between the microtubules and ciliary membrane with a radial peak position at 111±1 nm and a route width of 43±3 nm, and the other inside the axonemal lumen with a radial peak position at 0±1 nm and a route width of 98±3 nm (Figure 1 L-N). That the outer transport routes of both KIF3A and *α*-tubulin localize between the microtubules and ciliary membrane was expected as was the microtubule-proximal localization of the motor as compared to IFT proteins and cargo. We hypothesize that the inner route of both proteins corresponds to a passive diffusion route as we have previously observed in the cilium shaft (Luo et al 2017).

To map the passive diffusion route in the TZ, we localized the freely diffusing molecule GFP using Arl13b-mCherry as a ciliary marker. We found that GFP localizes at the center of the TZ with a peak position at 0±1 nm along the cilium radius and a route width of 70±1 nm (Figure 1 O-Q), which co-localizes well with the inner transport routes of the IFT motor and cargo. To demonstrate that transit of IFT motors and cargos in the lumen corresponds to passive diffusion, we examined the ATP dependence of this localization. We permeabilized NIH-3T3 cells with digitonin, which has been shown to selectively perforate the cellular membrane causing ATP to flow out of the cell (Breslow et al., 2013). This reduces ATP levels in the primary cilium by ∼90% inhibiting IFT (Figure 2 I) (Ye et al., 2013). We found that in permeabilized cells, the IFT motor KIF3A and the IFT cargo α-tubulin no longer occupied the outer transport route but continued to undergo transport through the luminal route (Figure 1 R-W). Thus, the luminal transport routes for α-tubulin and KIF3A are able to persist in an energy-independent environment via passive diffusion.

**Figure 2.**
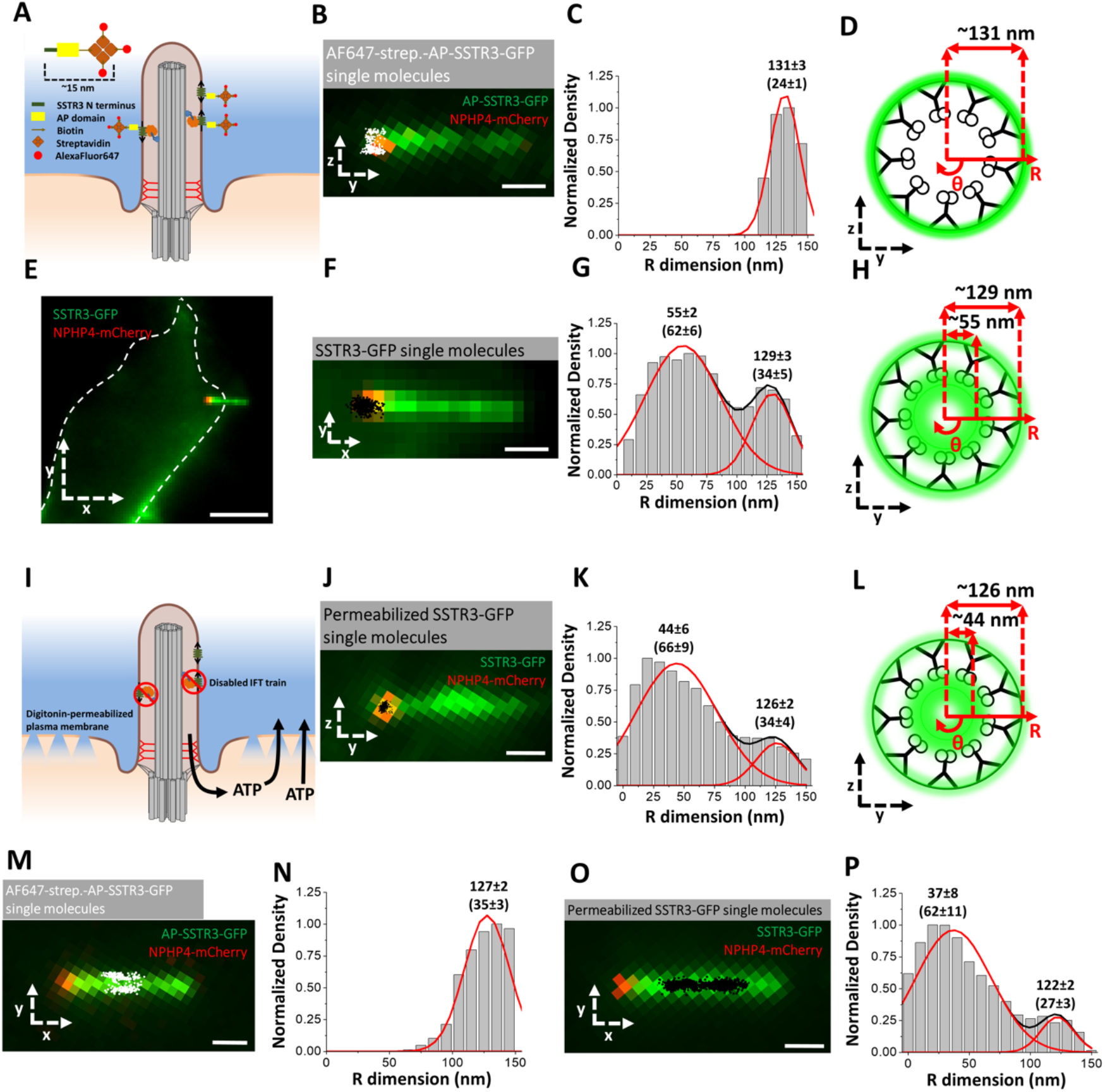
A previously uncharacterized transport route exists for membrane proteins inside the lumen of the primary cilium. **(A-D)** Imaging of externally-labeled AP-SSTR3-GFP. **A)** Schematic of the external labeling procedure. The extracellular SSTR3 N-terminus (green) is tagged with the AP (yellow) and biotinylated (arrow) for binding to AlexaFluor647 (red)-labeled streptavidin (orange). **B)** Image of primary cilium in live cells co-expressing AP-SSTR3-GFP (green) and NPHP4-mCherry (red) overlaid with 2D single-molecule Alexa-Fluor-647 externally labeled SSTR3 locations (white). Scale bar: 1 µm. **C)** Single-molecule AP-SSTR3-GFP locations in the TZ plotted along the R dimension in the cylindrical system. **D)** Spatial representation of the histogram in (C). **(E-H)** Imaging of SSTR3-GFP. **E)** Epifluorescence microscopy image of live cell co-expressing SSTR3-GFP (green) and NPHP4-mCherry (red) (scale bar: 5 µm). The dashed white line represents the cell border. **F)** Enlarged image of the primary cilium overlaid with single molecule SSTR3-GFP locations (black dots). Scale bar: 1 µm. Single-molecule data and epi-fluorescence images were rotated together clockwise, so that TZ is parallel to the x dimension to maintain consistency with subsequent data analyses. **G)** Single molecule SSTR3-GFP locations in the TZ along the R dimension in the cylindrical system. **H)** Spatial representation of the histogram in (G). **(I-L)** Imaging of SSTR3-GFP in permeabilized cells **I)** Schematic of the permeabilization procedure, resulting in ATP diffusion from the cell. **J)** Image of primary cilium in live cells co-expressing SSTR3-GFP (green) and NPHP4-mCherry (red) overlaid with 2D single-molecule SSTR3-GFP locations (white) in permeabilized cells. Scale bar: 1 µm. **K)** Single-molecule SSTR3-GFP locations in the TZ plotted along the R dimension in the cylindrical system. **L)** Spatial representation of the histogram in (K). **M) and N)** Same as (F) and (G) for AP-SSTR3-GFP in the ciliary shaft. **O) and P)** Same as (J) and (K) for SSTR3-GFP in the ciliary shaft in permeabilized cells.

### Tracking of TM proteins through the TZ shows their localizations at both the ciliary membrane and the ciliary lumen

To extend our technique to TM proteins, we investigated the transport route of an externally-labeled version of Somatostatin Receptor 3 (SSTR3), a G-protein coupled receptor that localizes to primary cilia and has critical functions in the hippocampus (Händel et al., 1999, Berbari et al., 2007, Einstein et al., 2010). We used an SSTR3 construct, AP-SSTR3-GFP, which can be externally labeled in live NIH-3T3 cells and marks the ciliary membrane (Howarth and Ting, 2008, Ye et al., 2013). The AP domain is an acceptor peptide attached to the N terminus of SSTR3 that, when expressed with the biotin ligase BirA, becomes biotinylated at the level of the endoplasmic reticulum. The biotinylated AP-SSTR3-GFP is trafficked to the ciliary membrane where it can be externally labeled using Alexa Fluor 647(AF647)-labeled streptavidin (Figure 2 A) (Ye et al., 2013, Ruba et al., 2018). As expected, the AF647-labeled AP-SSTR3-GFP localized near the TZ ciliary membrane with a peak position at 131±3 nm along the cilium radius and a route width of 24±1 nm (Figure 2 B-D), after taking into consideration the estimated length of the external label (Methods).

We also examine the localization of SSTR3-GFP at the TZ in live cells (Figure 2 E,F). Surprisingly, we found SSTR3-GFP occupies two transport paths through the TZ: one near the ciliary membrane with a peak position at 129±3 nm and a route width of 34±5 nm, and the other inside the TZ lumen with a peak position at 55±2 nm and a route width of 62±6 nm (Figure 2 G,H). Based on Monte Carlo simulations which estimate the localization error of the mean for the 3D density histograms (Figure S7 and S8 and Table 1), the inner and outer routes are spatially resolvable with localization precisions of 2.0 nm and 3.2 nm, respectively, due to the collection of a sufficient number of high-resolution single-molecule localizations. A comparison of the peak fitting areas determined that transiting molecules had a 73% frequency in the inner route and 27% in the outer route. The outer transport route for SSTR3-GFP likely corresponds to the ciliary membrane as it co-localizes with the externally-labeled AP-SSTR3-GFP (Figure 2 C,D). However, the inner transport route has not been visualized in previous measurements. It seems unlikely that the inner route reflects the GFP-labeled C-terminus of SSTR3 extending into the axonemal lumen as the 102 aa C-terminal domain of SSTR3 could extend a maximum of ∼44 nm (Ainavarapu et al., 2007), given an average length of ∼ 0.4 nm/aa and the 2-4 nm size of GFP, a distance that is not sufficient to bridge the ∼70-100 nm between the outer and inner transport routes. It also seems unlikely that free GFP is cleaved from SSTR3-GFP and contributes to the inner route localization as the inner route of SSTR3-GFP and the passive diffusion route of free GFP are well-separated by a distance of ∼ 50 nm. We conclude that our high-resolution 3D data in live cells may support a model for TM protein transport that utilizes the ciliary lumen to a greater extent than previously recognized.

Given that transit in the ciliary lumen occurs by passive diffusion for free GFP, IFT motors, and IFT cargos (Figure 1), we hypothesized that energy-independent transport may account for SSTR3 mobility in the lumenal route. To test this, we permeabilized the cells with digitonin and then performed SPEED microscopy. Although permeabilization did reduce the total frequency of single-molecule events (from 32.4±14.9 events/s to 10.1±1.6 events/s, Figure S2 C), the location of transiting SSTR3-GFP molecules was largely unaffected (Figure 2 I-L). The majority of SSTR3-GFP molecules continued to utilize the inner route (85% compared to 73% in unpermeabilized cells). The location of the SSTR3 inner transport route showed a slight shift towards the cilium center in permeabilized cells, with a peak position at 44±6 nm along the cilium radius and a route width of 66±9 nm (Figure 2 K,L). For the outer transport route, permeabilization and loss of ATP increased the immobile portion and decreased the directionally-moving SSTR3 molecules (determined via Mean Squared Displacement (MSD) analysis of single molecule trajectories), in agreement with others (Ye et al., 2013) (Figure S2 A,B).

To determine whether these SSTR3 transport routes occur in different regions of the primary cilium, we determined the 3D density map for SSTR3 at the basal body and the cilium shaft (Figure S3 A-G). At the basal body, using γ-tubulin-mCherry as a marker (Muresan et al., 1993), SSTR3 again had two transport routes: one near the outer basal body with a peak position at 120±4 nm and a route width of 40±6 nm and the other inside the lumen of the basal body with a peak position at 55±2 nm and a route width of 54±4 nm (Figure S3 H,K). These localizations suggest that the majority of SSTR3 likely enters and/or exits the transition zone through the lumen of the basal body rather than through the gaps in the 9+0 microtubules or through diffusion along the membrane. Along the cilium shaft, two transport routes were also detected for SSTR3-GFP (Figure S3 J,M) while only the outer route was detected for externally-labeled AP-STR3-GFP (Figure 2 M,N).

To determine whether this paradigm is similar for other TM proteins, we determined the 3D transport routes at the basal body, the TZ, and the cilium shaft for HTR6, a GPCR that binds serotonin and localizes to primary cilia, tagged with GFP on its C-terminus (Berbari et al., 2008). We found that HTR6 also occupies two transport routes, one in the lumen and one at the ciliary membrane, with similar locations as those of SSTR3. In comparison to SSTR3, an even larger proportion of HTR6 localized to the lumenal route (Figure S3 N-P). One possibility for this phenomenon may be that HTR6 undergoes higher turnover in its targeting to the primary cilium and thus spends more time in intraciliary and intracellular transport rather than embedded in the ciliary membrane.

### RAB8A, a regulator of ciliary vesicle targeting co-localizes and co-transports with SSTR3

The luminal localization of SSTR3 and HTR6 was surprising and suggests that vesicular trafficking may contribute to the luminal transport of TM proteins. Vesicular trafficking components such as Rabin8 and RAB8A promote docking of ciliary vesicles to the microtubule triplets that comprise the basal body (Nachury et al., 2007, Yoshimura et al., 2007). We thus hypothesized that RAB8A may mediate the transport of SSTR3. We measured the transport routes of GFP-RAB8A through the TZ and cilium shaft in live primary cilia. Interestingly, we found that RAB8A utilized two transport routes in both of these locations (Figure 3 A-F). The majority of transit events (87%) at the TZ localized to the luminal transport route with a mean position of 50±6 nm along the cilium radius and a route width of 75±11 nm while the remainder of events localized at an outer transport route with a mean position of 133±3 nm along the cilium radius and a route width of 40±7 nm (Figure 3 A-C).

**Figure 3.**
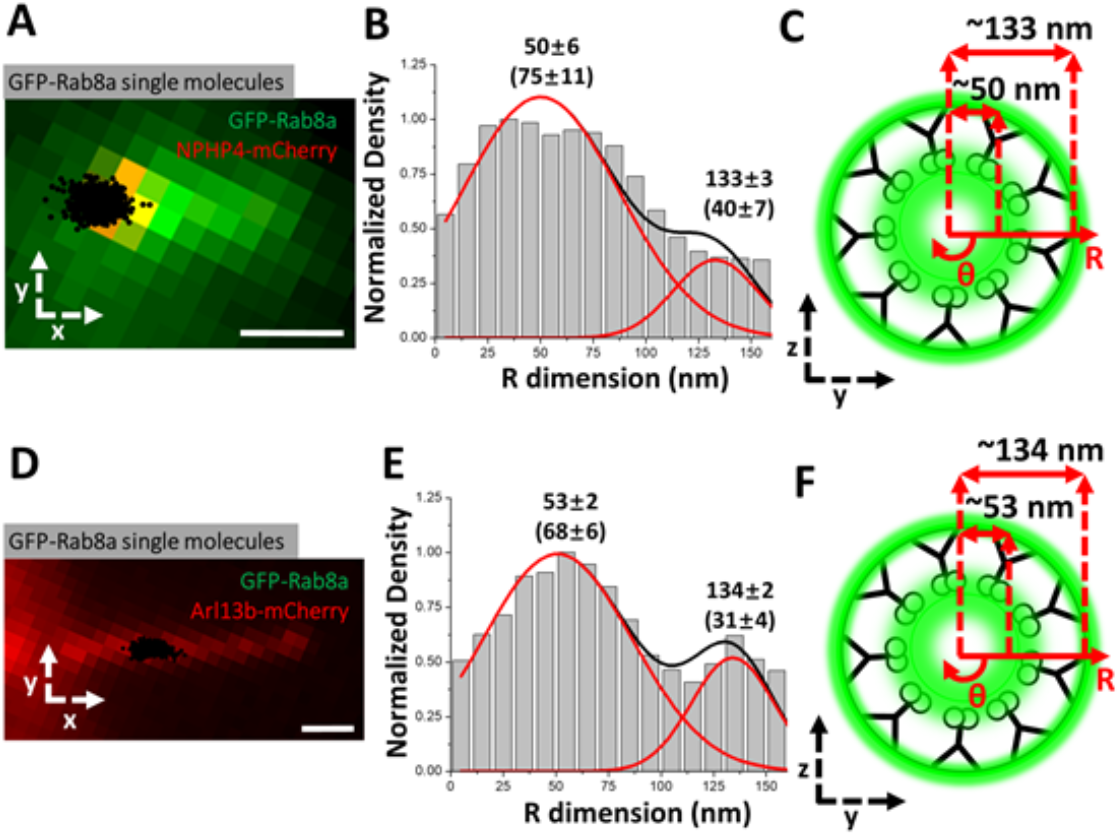
The transport routes for RAB8A are similar to those of SSTR3. **(A-C)** Imaging of GFP-RAB8A in the TZ. **A)** Epifluorescence microscopy image of GFP-RAB8A (green) and NPHP4-mCherry (red) overlaid with single-molecule GFP-RAB8A locations (black). Scale bar: 1 µm. **B)** Single-molecule GFP-RAB8A locations plotted along the R dimension. **C)** Spatial representation of (B). **(D-F)** Imaging of GFP-RAB8A in the cilium shaft. **D)** Epifluorescence microscopy image of GFP-RAB8A (green) and Arl13b-mCherry (red) overlaid with single-molecule GFP-RAB8A locations (black). Scale bar: 1 µm. **E)** Single-molecule GFP-RAB8A locations plotted along the R dimension. **F)** Spatial representation of (E).

RAB8A largely associates with membranes and has little turnover during the vesicle transport process in live cells (Grigoriev et al., 2011). The 3D transport routes for RAB8A, SSTR3 and HTR6 mapped by SPEED microscopy localize at the same position along the cilium radius that is outward from the passively-diffusing soluble molecules. We thus hypothesized that RAB8A may be tightly associated with vesicles bearing ciliary TM proteins, and causes them to hug the microtubule walls of the ciliary lumen. In this case, the single molecule trajectories of SSTR3 and RAB8A within primary cilia should be correlated. We utilized simultaneous dual-channel tracking of both SSTR3-mCherry and GFP-RAB8A in transfected NIH-3T3 cells and determined the angle difference between every step of each trajectory. In principle, an average angle difference close to 0° indicates a high degree of correlation between the trajectories of the proteins, whereas a wide distribution of angle differences indicates no correlation between the trajectory movements. We used measurements of the level of co-movement between positive and negative controls to generate a standard curve (Figure S4 M). The positive control experiments indicated that green and red dyes attached to 100-nm beads (Figure S4 C-F and Video 2) have a probability of 95% for co-movement, when “co-movement” was defined as the total angle differences of their trajectories that fell within ∼37° centered on the 0° bin (Figure S4 D-F). The negative control indicated that free fluorescein and JF561 dyes (Figure S4 G-J) have a probability of co-movement of ∼13% (Figure S4 I,J). By referencing the standard curve (Figure S4 M), we identified a probability of 24% co-movement translates to a 15% co-movement for SSTR3 and RAB8A trajectories in cilia above the negative control. Both the co-localization of the 3D transport and the degree of co-movement suggest that RAB8A may indeed by mediating the transport of SSTR3 (Figure S4 A,B,K,L).

### Golgicide A suppresses TM protein frequency equally in both routes and prevents ciliogenesis

As a second approach to test the role of vesicular trafficking in SSTR3 transport in the ciliary lumen, we treated cells with varying concentrations of Golgicide A (GCA), an inhibitor of Golgi export of TM proteins that works by inhibiting GBF1 from interacting with both Arf4 and the ciliary targeting sequence, a necessary step in sorting and export from the Golgi (Figure 4 A) (Wang et al., 2017, Wang et al., 2012). NIH-3T3 cells were serum starved in the absence or presence of GCA for 24 hr. GCA inhibited the length and ultimately the presence of primary cilia in a dose-dependent manner (Figure 4 B-D), indicating that membrane flux to primary cilia is an important component of ciliogenesis, particularly in relation to vesicle transport (Nachury et al., 2007). GCA treatment also reduced the frequency of single-molecule SSTR3-GFP events at the TZ in a dose-dependent manner (Figure 4 E). For the remaining events, the relative density and locations of the 3D transport routes were statistically unaffected (Figure 4 F-I), indicating that a similar mechanism of entry is utilized by both the inner and outer routes. It may be that the choice of which route to take is stochastic and based on the cross-sectional area of ciliary entry given that the space between the microtubules and the ciliary membrane accounts for ∼30% of the cross-sectional area while the ciliary lumen accounts for ∼70%. It thus appears that both the inner and outer SSTR3 transport routes are linked to vesicular transport mechanisms, with the inner route accommodating the bulk of SSTR3 transport in these experimental conditions.

**Figure 4.**
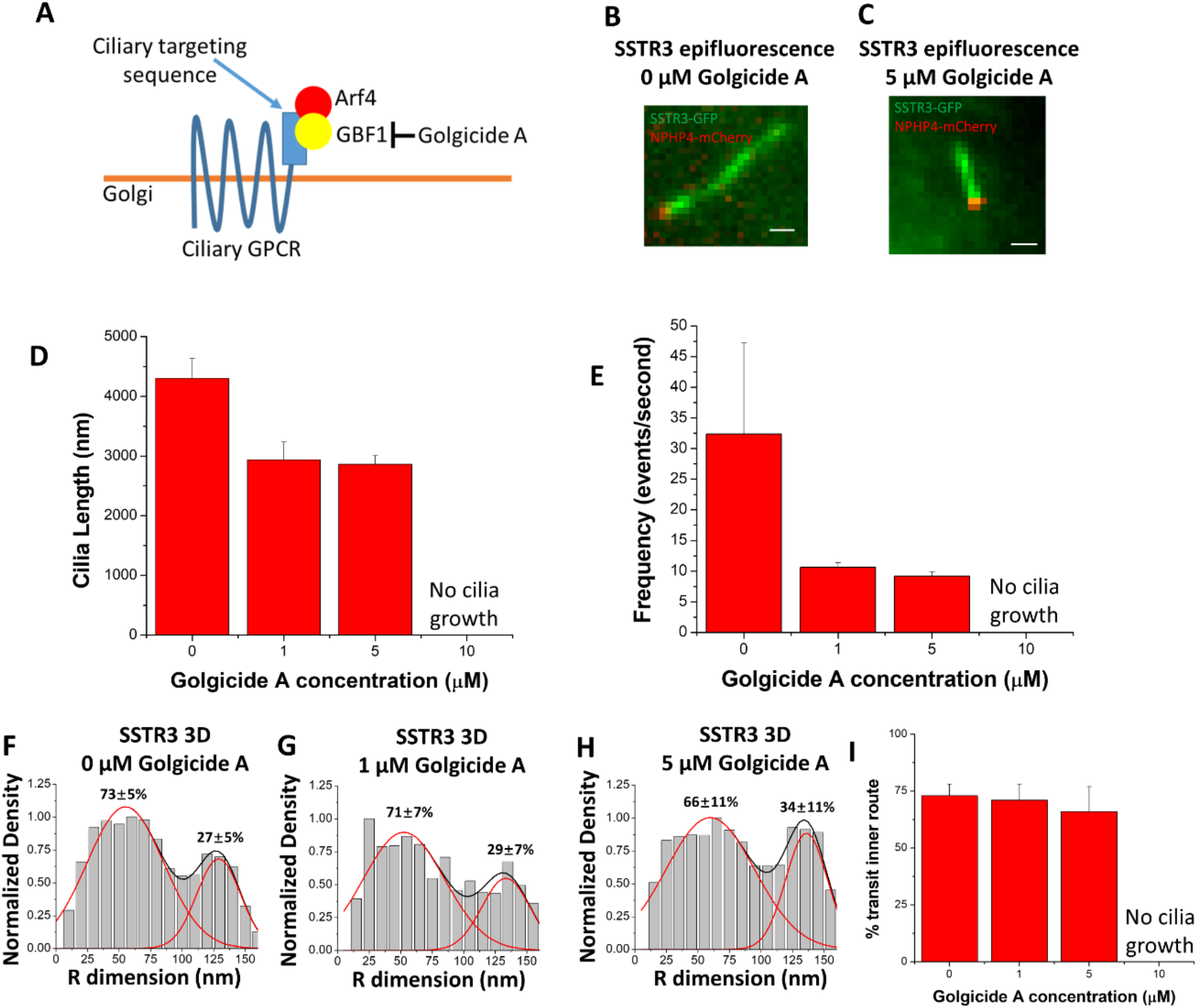
Golgicide A reduces SSTR3 frequency but does not alter transport routes. **A)** Schematic outlining Golgicide A’s mechanism of action. **B,C)** Epifluorescence images of SSTR3-GFP in NIH-3T3 cells treated with B) no Golgicide A or C) 5 µM Golgicide A for 24 hr. Scale bars: 1 µm. **D)** Bar graph showing cilia length vs. Golgicide A concentration. (0 µM n = 29, 1 µM n = 8, 5 µM n = 12, bar graph represents mean ± s.e.) **E)** Bar graph showing SSTR3 single molecule frequency vs. Golgicide A concentration (0 µM n = 5, 1 µM n = 3, 5 µM n = 4, bar graph represents mean ± s.e.) **F)-H)** 3D transport routes for SSTR3 in 0, 1, and 5 µM Golgicide A, respectively, plotted along the R dimension. **I)** Summary of percentage of SSTR3 transport in the inner transport route for 0, 1, 5, and 10 µM Golgicide A (statistics summarized in Table 1).

### The ciliary lumen is the preferred transport route for TM proteins during clathrin-mediated, signal-dependent internalization

Given the presence of both inner and outer routes for SSTR3 transit at the TZ, we wondered whether the relative utilization of these routes could be affected under conditions of active GPCR signaling. Previous work has shown that SSTR3 is actively removed from primary cilia after binding somatostatin (Green et al., 2016, Ye et al., 2018). We found that after stimulation with 10 µM somatostatin for 1 hr, the majority of SSTR3 single-molecule trajectories at the base of primary cilia displayed characteristics of both random diffusion and unclear movement patterns (Figure 5 D-I) while a minority displayed long, directional movement (Figure 5 A-C and Video 3). We then focused on every step of each trajectory and measured the average directionality of all the trajectories for each concentration of somatostatin in a range from 0-100 µM. As the concentration of somatostatin increased, the average directionality of trajectories shifted from 6% into the primary cilium to 16% out of the primary cilium (Figure 5 J), reflecting an overall removal of SSTR3 from the primary cilium, consistent with previous work (Ye et al., 2018). In concert with the shift in the average directionality of the trajectories, we found that the percent of transiting molecules in the lumenal transport route also increased until 100% of molecules utilized the inner route at 100 µM somatostatin (Figure 5 K-M,O).

**Figure 5.**
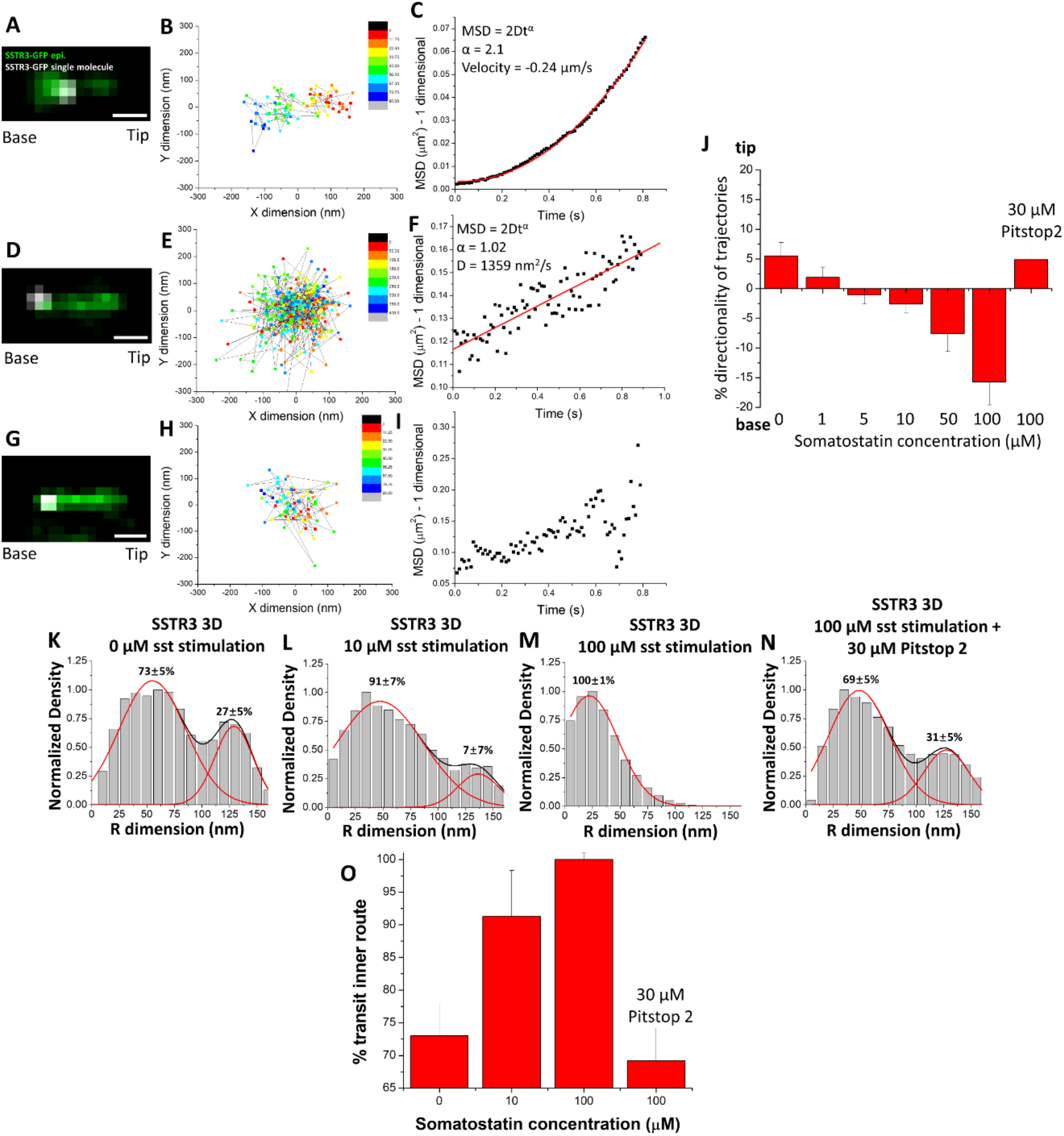
Somatostatin stimulation causes an increase in the transport of SSTR3 in the inner transport route. **A)** Epifluorescence image of SSTR3-GFP overlaid with single-molecule locations (white). Scale bar: 1 µm. **B)** 2D scatterplot of the Gaussian fitting results of the single molecule trajectory in (A). **C)** MSD vs. time plot of trajectory in (A) and (B). **D)-F) and G)-I)** Same as (A)-(C) showing trajectories with different types of movement patterns. **J)** Bar chart showing the average percent directionality of every trajectory step in different concentrations of somatostatin and Pitstop 2. (0 µM sst n = 489, 1 µM sst n = 908, 5 µM sst n = 889, 10 µM sst n = 1080, 50 µM sst n = 275, 100 µM sst n = 159, 100 µM sst/30 µM Pitstop 2 n = 427, bar graphs represent mean ± s.e.) **K)-N)** 3D transport routes in the TZ for SSTR3 in 0, 10, 100 µM somatostatin and 100 µM somatostatin with 30 µM Pitstop 2. **O)** Percentage of SSTR3-GFP molecules utilizing the inner transport route from (K)-(N). (statistics summarized in Table 1).

The mechanism for SSTR3 removal from primary cilia upon somatostatin stimulation has been shown to involve recruitment of β-arrestin to primary cilia, binding to SSTR3, and promotion of clathrin-coated endocytosis (Green et al., 2016, Oakley et al., 1999). To investigate the role of clathrin-coated endocytosis in SSTR3 removal from primary cilia, we measured the directionality and 3D transport route of SSTR3 at 100 µM somatostatin following a 1 hour pre-incubation with 30 µM Pitstop 2, a potent clathrin-coated endocytosis inhibitor (von Kleist et al., 2011). Pitstop 2 treatment completely blocked the dose-dependent reversal of SSTR3 directionality during somatostatin stimulation (Figure 5 J) and prevented the shift of transiting SSTR3 to the luminal transport route (Figure 5 N,O) suggesting that clathrin-coated endocytosis plays a role in facilitating movement through the inner transport route.

### The ciliary lumen is the preferred transport route for TM proteins during ciliogenesis

As a second test of whether the relative utilization of the inner and outer transport routes could be affected by cellular conditions, we examined the transit of SSTR3 and HTR6 molecules during different stages of ciliary growth. Due to the non-linear growth kinetics of primary cilia (Figure 6 A), we selected time points of 1, 3, 6, 12, and 24 hours following serum starvation. For both SSTR3 and HTR6, the majority of transit events at the TZ occurred via the luminal transport route early in ciliary growth and shifted to include more outer transit events as the primary cilia matured (Figure 6 B-L). We used the Pearson correlation coefficient to correlate these changes with values close to +1 or −1 indicating high degrees of positive and negative correlation, respectively, and values close to 0 indicating no correlation. Changes in percent transit in the inner route and ciliary length had strong negative correlation at −0.90 and −0.83 for SSTR3 and HTR6, respectively (Figure 6 M), suggesting that these phenomena may arise from the same biological process that regulates ciliogenesis. The percent transit in the inner route showed no correlation with the frequency of single-molecule events (Pearson coefficients of 0.13 and 0.15 for SSTR3 and HTR6, respectively, Figure S5). These results suggest that, unlike structural and IFT proteins in flagellum elongation, TM proteins may not be transported to the growing cilium at a higher rate earlier in ciliogenesis. Thus, the changes in percent transit in the central transport route may be due to other factors, such as structural and selective competency of the transition zone which forms early in ciliogenesis (Williams et al., 2011).

**Figure 6.**
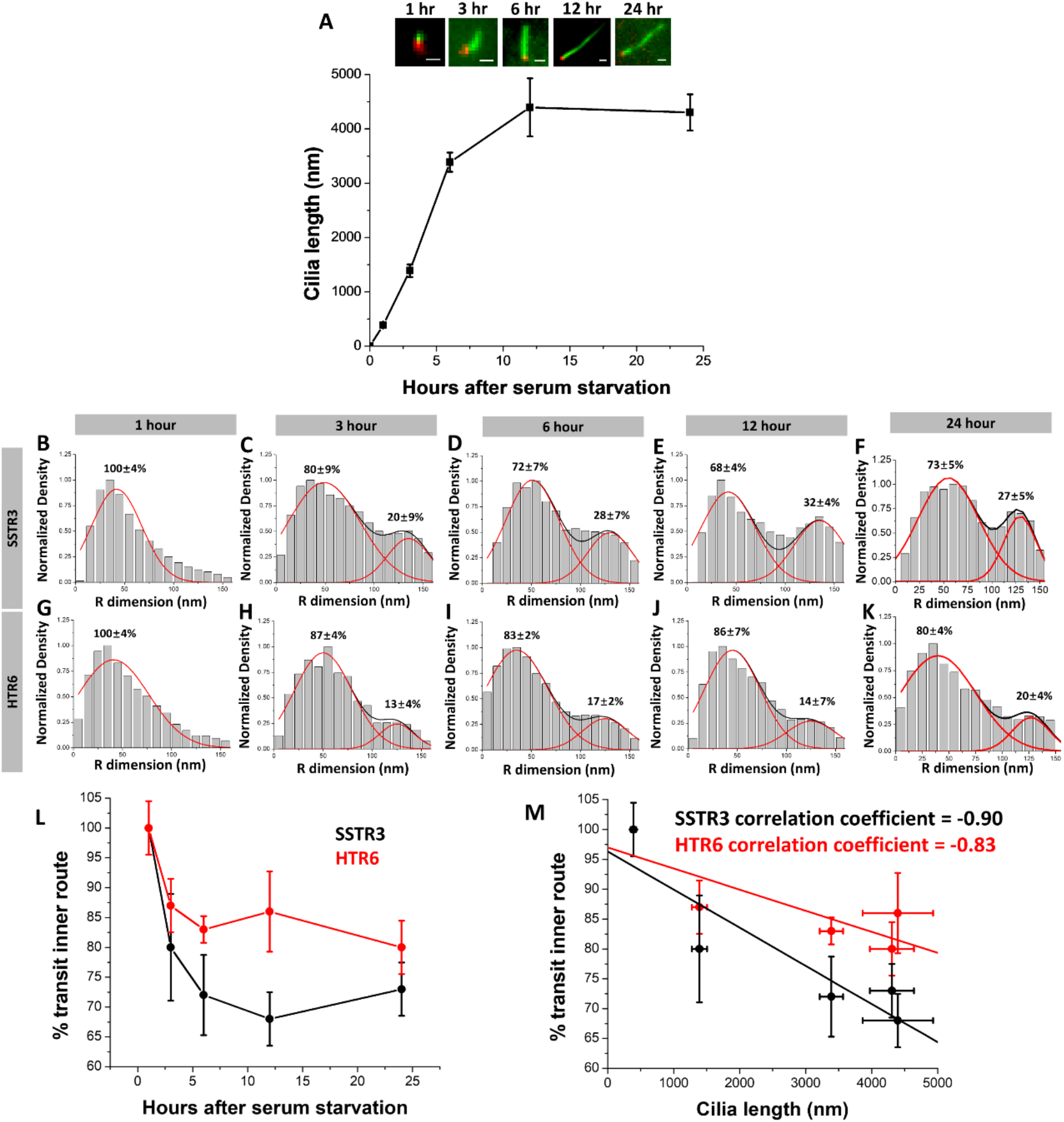
SSTR3 and HTR6 use the inner transport route at a higher frequency at earlier times of ciliogenesis. **A)** Growth curve of primary cilia following serum starvation. Insets: green = SSTR3-GFP, red = NPHP4-mCherry. **B)-F)** 3D transport routes for SSTR3 following 1, 3, 6, 12, and 24 hours of serum starvation. **G)-K)** Same as (B)-(F) except with HTR6. **L)** Summary of (B)-(K). **M)** Graph of percent transit in inner route vs. average cilia length with Pearson’s correlation coefficient for both SSTR3 and HTR6.

### The ciliary lumen accommodates similar transport routes in Chlamydomonas reinhardtii flagella

The utilization of an inner transport route by a TM protein in the TZ of mammalian primary cilia was surprising and we sought to determine whether such events occur in other cilia. Therefore, we performed single molecule tracking (Video 4) and determined the 3D transport routes for several classes of proteins in *Chlamydomonas reinhardtii* flagella: the IFT cargo β-tubulin (Craft et al., 2015), the TM protein PKD2 (Huang et al., 2007), the IFT protein IFT54 (Wingfield et al., 2017), and the kinesin-2 motor subunit KAP (Mueller et al., 2005). Overall, the results were quite similar to mammalian primary cilia in terms of transport routes observed for each protein. β-tubulin, PKD2, and KAP all had inner and outer transport routes in the shaft of the *Chlamydomonas* flagellum (Figure 7 A-J) that roughly correlated to the same transport routes for α-tubulin, SSTR3, and KIF3A in primary cilia (Figure 1). One difference between the systems was a radial shift in the inner routes for β-tubulin and KAP by 11 nm and 29 nm, respectively, perhaps due to the presence of the central pair of microtubules in the motile *Chlamydomonas* flagellum (Czarnecki and Shah, 2012). A second difference was a shift of the IFT54 transport route to 79 nm along the radius of flagella which is ∼25 nm more central compared to IFT20 and IFT43 in primary cilia. This discrepancy may be due to slightly different locations of the axoneme in flagella and cilia and/or different placement of IFT54 within the IFT particle compared to IFT20 and IFT43. Taken together, the general location of the 3D transport routes appears to be similar in the motile flagellum of *Chlamydomonas* and the primary cilium of mammalian cells.

**Figure 7.**
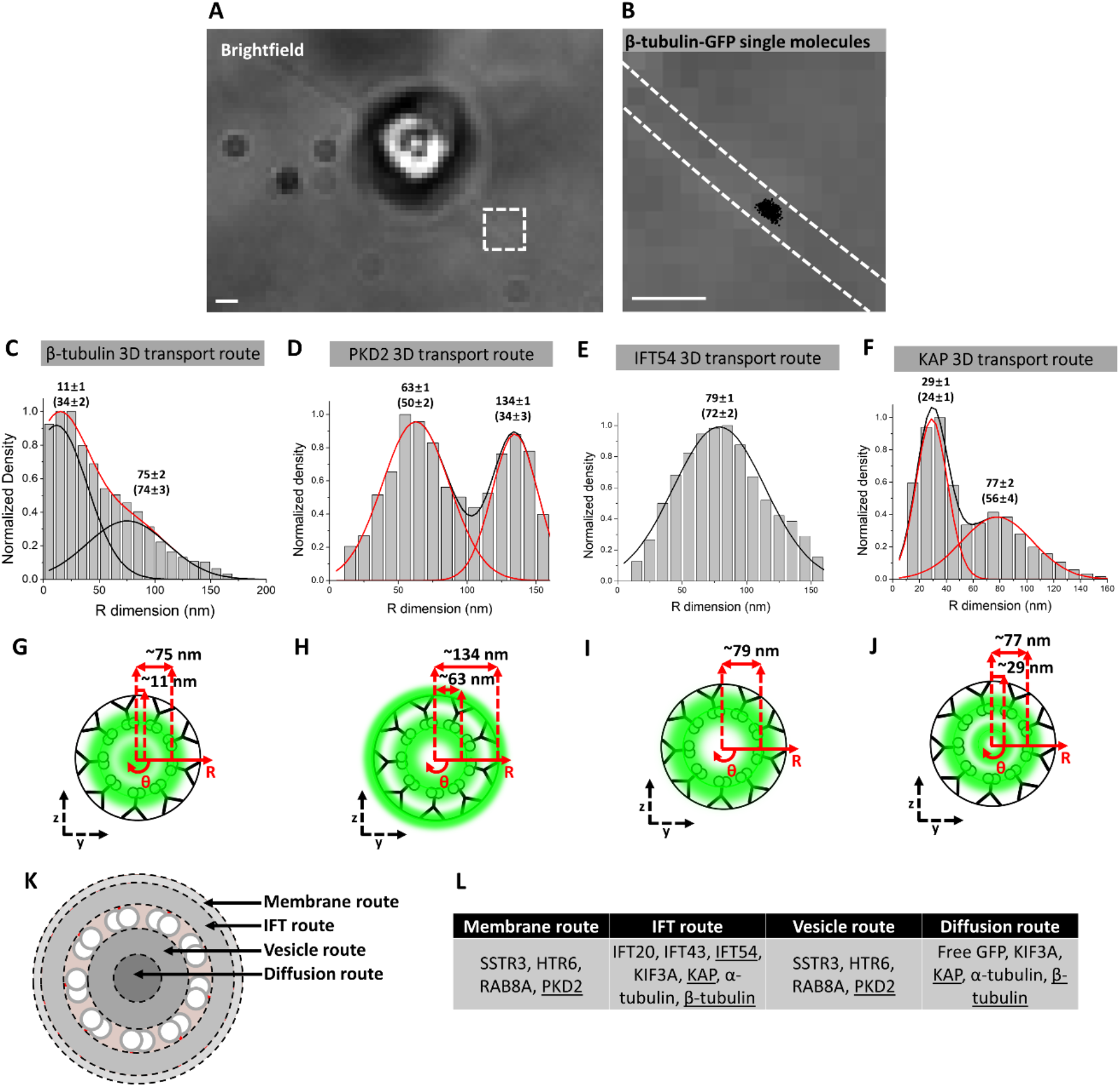
Transport routes in Chlamydomonas reinhardtii flagella closely reflect those in primary cilia. **A)** Brightfield image of *Chlamydomonas reinhardtii* stably expressing β-tubulin-GFP. Red dashed box shows the region of flagellum selected for single-molecule imaging. Scale bar: 1 µm. **B)** Brightfield image from boxed region in (A) with dashed white lines indicating location of flagellum and superimposed with single-molecule localizations (black dots). Scale bar: 1 µm. **C)-F)** 3D transport routes for β-tubulin, PKD2, IFT54, and KAP, respectively, in the shaft of the flagellum. **G)-J)** Spatial representations of (C)-(F), respectively. **K)** Summary of various transport routes observed in a cross-section view of a primary cilium. **L)** Table indicating which proteins utilize the transport routes depicted in (K). Underlined proteins indicate transport routes determined in *Chlamydomonas reinhardtii*.

## Discussion

### Vesicles as transmembrane protein carriers in the ciliary lumen

Our results show that the lumen of primary cilia is a prominent pathway for the transport of TM proteins in two model systems and that its relative usage can be regulated by a variety of ciliary states. The ciliary lumen also accommodates the transport of passively diffusing soluble molecules including α-tubulin, kinesin-2, and free GFP. We note that the axonemal lumen transport route is offset by ∼50 nm from the passive diffusion route in the central axis (Figure 7 K). That vesicle transport is involved in the inner route is supported by 1) co-localization with Rab8A, 2) the reduction in transport in response to GCA treatment, and 3) sensitivity to Pitstop 2 treatment. A summary of the transport routes and the proteins that traverse these routes can be found in Figure 7 K,L.

Our results characterize the ciliary lumen as a viable transport route for vesicles, a finding that has been entertained in the field based on EM studies of chondrocytes and rod photoreceptors (Jensen et al., 2004, Chuang et al., 2015). Recent work has also visualized vesicles in the amphid cilia of *C. elegans*, and vesicles accumulated in worms null for various IFT transport proteins (Li et al., 2019). A criticism of the EM work has been the propensity of the EM chemical fixation process to form vesicle-like bubbles. EM is also limited in its ability to resolve low-contrast structures like vesicles unless vesicular proteins are specifically and densely labeled by immunogold labeling (Nachury et al., 2010). In fact, early characterization studies of SSTR3 localization in the hippocampus showed dense immunogold labeling along the entire width of primary cilia (Händel et al., 1999) and electron-dense particulate collects in the empty space of primary cilia (Rogowski et al, 2013). Thus, EM would be ideal for observing vesicle transport of TM proteins in primary cilia if not for its main limitations. Therefore, we opted for a live-cell, super-resolution, and fluorescence-based approach to study this question.

We validated the ability of our SPEED microscopy and 2D-to-3D transformation algorithm to accurately reconstruct known details of protein transport in primary cilia using IFT components and the AP-SSTR3-GFP construct. These results elucidate the transport routes and mechanisms for TM and other proteins in primary cilia. We also examined the effects of single molecule localization precision (Figure S8), number of single molecule localizations (Figure S9), labeling efficiency (Figure S10), and distortion (Figure S11) on the precision of the final 3D transport routes via Monte Carlo simulation (Figure S7) for every transport route presented in this paper. Each transport route conforms to the precision standards of the biological claims we make according to simulation and the results are summarized in Table 1. Furthermore, we emphasize the reasonable consistency of the ciliary ultrastructure and the experimental reproducibility of each transport route by showing the 3D transport route histograms for SSTR3 in 6 different cilia and quantify the error (Figure S12).

Even though the lumen of the TZ is a relatively open and clear channel compared to the surrounding structural regions and likely free from the cartwheel structure that characterizes non-ciliary procentrioles (Alvey, 1986, Hoyer-Fender, 2012), an important consideration is how vesicles may pass through the basal body, TZ, and cilium shaft. First, there may be some vesicle distortion that occurs to allow vesicle passage into and along primary cilia. Indeed, the recycling endosome, a waypoint for TM proteins sent to the primary cilia, has a convoluted membrane structure and its vesicles are highly non-uniform (Goldenring, 2015). Second, there appears to be a requirement for proteins to mediate the transport of vesicles past the diffusion barrier into primary cilia. For example, RAB8A is responsible for mediating the entry into the connecting cilium of frog photoreceptors (Moritz et al., 2001) and we show that it occupies the same transport routes as SSTR3 *in vivo*. Third, small vesicles carrying TM proteins may be able to pass through the diffusion barrier, albeit at a low frequency due to being above the widely accepted barrier limit (Kee et al., 2012, Breslow et al., 2013, Calvert et al., 2010, Lin et al., 2013).

Previous models have suggested that the fusion of TM protein-containing vesicles likely occurs outside the primary cilia, at the periciliary base, or at the TZ ciliary membrane (Nachury et al., 2010). Overall, our results do not refute these findings. Indeed, RAB8A possessed transport routes in the lumen and near the ciliary membrane. In addition, inhibition of TM protein vesicle export from Golgi with GCA showed reduction in both the inner and outer transport routes. This suggests that vesicle transport is a component of transiting both TZ locations. Our work aims to parse out the usage of both of these routes during the different ciliary states.

### The ciliary lumen as a specialized signaling transport route

Several groups have shown via fluorescence microscopy that markers for clathrin-coated pits localize to the ciliary pocket (Molla-Herman et al., 2010) and others find that β-arrestin is recruited to primary cilia upon SSTR3 stimulation and has a causal role in its removal (Green et al., 2016). Based on this evidence, it appears that β-arrestin is recruited to primary cilia following SSTR3 stimulation, attenuates SSTR3’s signaling, and facilitates transport by IFT out of primary cilia where it promotes interaction with endocytic machinery. Our data extend this model by suggesting that the ciliary membrane possesses endocytic capabilities and that the internalized receptors can be transported through the axonemal lumen.

The primary cilium is a specialized signaling organelle whose evolutionary progenitor is the motile flagellum (Mitchell., 2007). Over time and with selection pressure against motility in eukaryotic cells, it appears that proto-primary cilia lost the central pair of microtubules and associated motility proteins. This loss of motility function may have paved the way for an increase in the signaling capabilities by reducing the geometric constraints of the flagellum. Here, we show that TM protein transport at various stages of the primary cilium lifecycle – growth, steady-state, and signaling states – can occur, at least in part, through the axonemal lumen. In future work, probing the differences between flagella and primary cilia and incorporating other well-known ciliary protein transport components, such as the BBSome, will be essential in developing a molecular view of the various ciliopathies.

## Acknowledgements

The project was supported by grants from the National Institutes of Health (NIH GM097037, GM116204 and GM122552 to W.Y.). We also acknowledge Dr. Joel Rosenbaum (Yale University) for critical comments on the manuscript.

## Author Contributions

In this manuscript, A.R., W.L., and W.Y. designed experiments; A.R., W.L. and D.T. prepared plasmids and established cell lines; A.R., W.L. and W.Y. performed single-molecule tracking, super-resolution SPEED microscopy, and wide field microscopy experiments; A.R., W.L. and W.Y. conducted data analysis; A.R., W.L., K.J.V. and W.Y. wrote the manuscript.

## References

Ainavarapu, Sri Rama Koti, et al. “Contour length and refolding rate of a small protein controlled by engineered disulfide bonds.” Biophysical journal 92.1 (2007): 225–233.

Alvey, P. L. “Do adult centrioles contain cartwheels and lie at right angles to each other?.” Cell biology international reports 10.8 (1986): 589–598.

Anholt, Robert RH, et al. “Transduction proteins of olfactory receptor cells: identification of guanine nucleotide binding proteins and protein kinase C.” Biochemistry 26.3 (1987): 788–795.

Anvarian, Zeinab, et al. “Cellular signalling by primary cilia in development, organ function and disease.” Nature Reviews Nephrology (2019): 1.

Arts, Heleen H., et al. “C14ORF179 encoding IFT43 is mutated in Sensenbrenner syndrome.” Journal of medical genetics 48.6 (2011): 390–395.

Awata, Junya, et al. “Nephrocystin-4 controls ciliary trafficking of membrane and large soluble proteins at the transition zone.” J Cell Sci (2014): jcs-155275.

Bangs, Fiona, and Kathryn V. Anderson. “Primary cilia and mammalian hedgehog signaling.” Cold Spring Harbor perspectives in biology 9.5 (2017): a028175.

Barzi, Mercedes, et al. “Sonic Hedgehog-induced proliferation requires specific Gα inhibitory proteins.” Journal of Biological Chemistry 286.10 (2011): 8067–8074.

Berbari, Nicolas F., et al. “Hippocampal neurons possess primary cilia in culture.” Journal of neuroscience research 85.5 (2007): 1095–1100.

Berbari, Nicolas F., et al. “Identification of ciliary localization sequences within the third intracellular loop of G protein-coupled receptors.” Molecular biology of the cell 19.4 (2008): 1540–1547.

Breslow, David K., et al. “A CRISPR-based screen for Hedgehog signaling provides insights into ciliary function and ciliopathies.” Nature genetics 50.3 (2018): 460.

Breslow, David K., et al. “An in vitro assay for entry into cilia reveals unique properties of the soluble diffusion barrier.” J Cell Biol 203.1 (2013): 129–147.

Breunig, Joshua J., et al. “Primary cilia regulate hippocampal neurogenesis by mediating sonic hedgehog signaling.” Proceedings of the National Academy of Sciences 105.35 (2008): 13127–13132.

Brooks, Celine, et al. “Farnesylation of the Transducin G protein gamma subunit is a prerequisite for its ciliary targeting in rod photoreceptors.” Frontiers in molecular neuroscience 11 (2018): 16.

Burke, Michael C., et al. “Chibby promotes ciliary vesicle formation and basal body docking during airway cell differentiation.” J Cell Biol 207.1 (2014): 123–137.

Calvert, Peter D., William E. Schiesser, and Edward N. Pugh. “Diffusion of a soluble protein, photoactivatable GFP, through a sensory cilium.” The Journal of general physiology 135.3 (2010): 173–196.

Chuang, Jen-Zen, Ya-Chu Hsu, and Ching-Hwa Sung. “Ultrastructural visualization of trans-ciliary rhodopsin cargoes in mammalian rods.” Cilia 4.1 (2015): 4.

Corbit, Kevin C., et al. “Vertebrate Smoothened functions at the primary cilium.” Nature 437.7061 (2005): 1018.

Craft, Julie M., et al. “Tubulin transport by IFT is upregulated during ciliary growth by a cilium-autonomous mechanism.” J Cell Biol 208.2 (2015): 223–237.

Craige, Branch, et al. “CEP290 tethers flagellar transition zone microtubules to the membrane and regulates flagellar protein content.” The Journal of cell biology 190.5 (2010): 927–940.

Czarnecki, Peter G., and Jagesh V. Shah. “The ciliary transition zone: from morphology and molecules to medicine.” Trends in cell biology 22.4 (2012): 201–210.

Deschout, Hendrik, Kristiaan Neyts, and Kevin Braeckmans. “The influence of movement on the localization precision of sub-resolution particles in fluorescence microscopy.” Journal of biophotonics 5.1 (2012): 97–109.

Dirksen, Ellen Roter, and Ilona Staprans. “Tubulin synthesis during ciliogenesis in the mouse oviduct.” Developmental biology 46.1 (1975): 1–13.

Dosztányi, Zsuzsanna, et al. “IUPred: web server for the prediction of intrinsically unstructured regions of proteins based on estimated energy content.” Bioinformatics 21.16 (2005): 3433–3434.

Einstein, Emily B., et al. “Somatostatin signaling in neuronal cilia is criticalfor object recognition memory.” Journal of Neuroscience 30.12 (2010): 4306–4314.

Engelke, Martin F., et al. “Acute inhibition of heterotrimeric kinesin-2 function reveals mechanisms of intraflagellar transport in mammalian cilia.” Current Biology (2019).

Erickson, Harold P. “Size and shape of protein molecules at the nanometer level determined by sedimentation, gel filtration, and electron microscopy.” Biological procedures online 11.1 (2009): 32.

Follit, John A., et al. “The intraflagellar transport protein IFT20 is associated with the Golgi complex and is required for cilia assembly.” Molecular biology of the cell 17.9 (2006): 3781–3792.

Gaffield, Michael A., Silvio O. Rizzoli, and William J. Betz. “Mobility of synaptic vesicles in different pools in resting and stimulated frog motor nerve terminals.” Neuron 51.3 (2006): 317–325.

Gerdes, J. M., and N. Katsanis. “Microtubule transport defects in neurological and ciliary disease.” Cellular and Molecular Life Sciences CMLS 62.14 (2005): 1556–1570.

Goldenring, James R. “Recycling endosomes.” Current opinion in cell biology 35 (2015): 117–122.

Green, Jill A., et al. “Recruitment of β-arrestin into neuronal cilia modulates somatostatin receptor subtype 3 ciliary localization.” Molecular and cellular biology 36.1 (2016): 223–235.

Grigoriev, Ilya, et al. “Rab6, Rab8, and MICAL3 cooperate in controlling docking and fusion of exocytotic carriers.” Current Biology 21.11 (2011): 967–974.

Han, Weiping, et al. “Neuropeptide release by efficient recruitment of diffusing cytoplasmic secretory vesicles.” Proceedings of the National Academy of Sciences 96.25 (1999): 14577–14582.

Händel, M., et al. “Selective targeting of somatostatin receptor 3 to neuronal cilia.” Neuroscience 89.3 (1999): 909–926.

Hao, Limin, et al. “Intraflagellar transport delivers tubulin isotypes to sensory cilium middle and distal segments.” Nature cell biology 13.7 (2011): 790.

He, Xuelian, et al. “The G protein α subunit Gα s is a tumor suppressor in Sonic hedgehog− driven medulloblastoma.” Nature medicine 20.9 (2014): 1035.

Higginbotham, Holden, et al. “Arl13b in primary cilia regulates the migration and placement of interneurons in the developing cerebral cortex.” Developmental cell 23.5 (2012): 925–938.

Hilgendorf, Keren I., Carl T. Johnson, and Peter K. Jackson. “The primary cilium as a cellular receiver: organizing ciliary GPCR signaling.” Current opinion in cell biology 39 (2016): 84–92.

Hirano, Tomoaki, Yohei Katoh, and Kazuhisa Nakayama. “Intraflagellar transport-A complex mediates ciliary entry and retrograde trafficking of ciliary G protein–coupled receptors.” Molecular biology of the cell 28.3 (2017): 429–439.

Horn, Meryl E., and Roger A. Nicoll. “Somatostatin and parvalbumin inhibitory synapses onto hippocampal pyramidal neurons are regulated by distinct mechanisms.” Proceedings of the National Academy of Sciences 115.3 (2018): 589–594.

Howarth, Mark, and Alice Y. Ting. “Imaging proteins in live mammalian cells with biotin ligase and monovalent streptavidin.” Nature protocols 3.3 (2008): 534.

Hoyer-Fender, Sigrid. “Primary and motile cilia: their ultrastructure and ciliogenesis.” Cilia and Nervous System Development and Function. Springer, Dordrecht, 2013. 1–53.

Huang, Kaiyao, et al. “Function and dynamics of PKD2 in Chlamydomonas reinhardtii flagella.” The Journal of cell biology 179.3 (2007): 501–514.

Hunnicutt, Gary R., Maria G. Kosfiszer, and William J. Snell. “Cell body and flagellar agglutinins in Chlamydomonas reinhardtii: the cell body plasma membrane is a reservoir for agglutinins whose migration to the flagella is regulated by a functional barrier.” The Journal of Cell Biology 111.4 (1990): 1605–1616.

Ishikawa, Hiroaki, et al. “Proteomic analysis of mammalian primary cilia.” Current Biology 22.5 (2012): 414–419.

Jana, Swadhin Chandra, et al. “Differential regulation of transition zone and centriole proteins contributes to ciliary base diversity.” Nature cell biology 20.8 (2018): 928.

Jenkins, Paul M., et al. “Ciliary targeting of olfactory CNG channels requires the CNGB1b subunit and the kinesin-2 motor protein, KIF17.” Current biology 16.12 (2006): 1211–1216.

Jensen, Victor L., and Michel R. Leroux. “Gates for soluble and membrane proteins, and two trafficking systems (IFT and LIFT), establish a dynamic ciliary signaling compartment.” Current opinion in cell biology 47 (2017): 83–91.

Jensen, C. G., et al. “Ultrastructural, tomographic and confocal imaging of the chondrocyte primary cilium in situ.” Cell biology international 28.2 (2004): 101–110.

Jones, Chonnettia, et al. “Ciliary proteins link basal body polarization to planar cell polarity regulation.” Nature genetics 40.1 (2008): 69.

Kaneshiro, E. S. “Lipids of Paramecium.” Journal of lipid research 28.11 (1987): 1241-1258.

Kee, Hooi Lynn, et al. “A size-exclusion permeability barrier and nucleoporins characterize a ciliary pore complex that regulates transport into cilia.” Nature cell biology 14.4 (2012): 431.

Kozminski, Keith G., Peter L. Beech, and Joel L. Rosenbaum. “The Chlamydomonas kinesin-like protein FLA10 is involved in motility associated with the flagellar membrane.” The Journal of cell biology 131.6 (1995): 1517–1527.

Kyoung, Minjoung, and Erin D. Sheets. “Vesicle diffusion close to a membrane: intermembrane interactions measured with fluorescence correlation spectroscopy.” Biophysical journal 95.12 (2008): 5789–5797.

Lerea, Connie L., et al. “Identification of specific transducin alpha subunits in retinal rod and cone photoreceptors.” Science 234.4772 (1986): 77-80.

Li, Ming, et al. “Ciliopathy-associated proteins are involved in vesicle distribution in sensory cilia.” Journal of genetics and genomics 46 (2019): 269–271.

Lin, Yu-Chun, et al. “Chemically inducible diffusion trap at cilia reveals molecular sieve–like barrier.” Nature chemical biology9.7 (2013): 437.

Luo, Wangxi, et al. “Axonemal lumen dominates cytosolic protein diffusion inside the primary cilium.” Scientific reports7.1 (2017): 15793.

Ma, Jiong, and Weidong Yang. “Three-dimensional distribution of transient interactions in the nuclear pore complex obtained from single-molecule snapshots.” Proceedings of the National Academy of Sciences 107.16 (2010): 7305–7310.

Ma, Jiong, Joseph M. Kelich, and Weidong Yang. “SPEED microscopy and its application in nucleocytoplasmic transport.” The Nuclear Envelope: Methods and Protocols (2016): 503–518.

Marshall, Wallace F., and Shigenori Nonaka. “Cilia: tuning in to the cell’s antenna.” Current Biology 16.15 (2006): R604–R614.

Mitchell, David R. “The evolution of eukaryotic cilia and flagella as motile and sensory organelles.” Eukaryotic Membranes and Cytoskeleton. Springer, New York, NY, 2007. 130–140.

Molla-Herman, Anahi, et al. “The ciliary pocket: an endocytic membrane domain at the base of primary and motile cilia.” Journal of cell science (2010): jcs-059519.

Moritz, Orson L., et al. “Mutant rab8 Impairs docking and fusion of rhodopsin-bearing post-Golgi membranes and causes cell death of transgenic Xenopus rods.” Molecular biology of the cell 12.8 (2001): 2341–2351.

Mortensen, Kim I., et al. “Optimized localization analysis for single-molecule tracking and super-resolution microscopy.” Nature methods 7.5 (2010): 377.

Mueller, Joshua, et al. “The FLA3 KAP subunit is required for localization of kinesin-2 to the site of flagellar assembly and processive anterograde intraflagellar transport.” Molecular biology of the cell 16.3 (2005): 1341–1354.

Muresan, Virgil, Harish C. Joshi, and Joseph C. Besharse. “Gamma-tubulin in differentiated cell types: localization in the vicinity of basal bodies in retinal photoreceptors and ciliated epithelia.” Journal of Cell Science 104.4 (1993): 1229–1237.

Nachury, Maxence V., E. Scott Seeley, and Hua Jin. “Trafficking to the ciliary membrane: how to get across the periciliary diffusion barrier?.” Annual review of cell and developmental biology 26 (2010): 59–87.

Nachury, Maxence V., et al. “A core complex of BBS proteins cooperates with the GTPase Rab8 to promote ciliary membrane biogenesis.” Cell 129.6 (2007): 1201–1213.

Nager, Andrew R., et al. “An actin network dispatches ciliary GPCRs into extracellular vesicles to modulate signaling.” Cell 168.1 (2017): 252–263.

Nachury, Maxence V., and David U. Mick. “Establishing and regulating the composition of cilia for signal transduction.” Nature Reviews Molecular Cell Biology (2019): 1.

Nair, K. Saidas, et al. “Light-dependent redistribution of arrestin in vertebrate rods is an energy-independent process governed by protein-protein interactions.” Neuron 46.4 (2005): 555–567.

Najafi, Mehdi, Nycole A. Maza, and Peter D. Calvert. “Steric volume exclusion sets soluble protein concentrations in photoreceptor sensory cilia.” Proceedings of the National Academy of Sciences 109.1 (2012): 203–208.

Nauli, Surya M., et al. “Polycystins 1 and 2 mediate mechanosensation in the primary cilium of kidney cells.” Nature genetics 33.2 (2003): 129.

Oakley, Robert H., et al. “Association of β-arrestin with G protein-coupled receptors during clathrin-mediated endocytosis dictates the profile of receptor resensitization.” Journal of Biological Chemistry 274.45 (1999): 32248–32257.

O’Hagan, Robert, et al. “Glutamylation regulates transport, specializes function, and sculpts the structure of cilia.” Current Biology 27.22 (2017): 3430–3441.

Ostrowski, Lawrence E., et al. “A proteomic analysis of human cilia: identification of novel components.” Molecular & Cellular Proteomics 1.6 (2002): 451–465.

Ott, Carolyn, and Jennifer Lippincott- Schwartz. “Visualization of live primary cilia dynamics using fluorescence microscopy.” Current protocols in cell biology 57.1 (2012): 4–26.

Papermaster, David S., B. G. Schneider, and J. C. Besharse. “Vesicular transport of newly synthesized opsin from the Golgi apparatus toward the rod outer segment. Ultrastructural immunocytochemical and autoradiographic evidence in Xenopus retinas.” Investigative ophthalmology & visual science 26.10 (1985): 1386–1404.

Pazour, Gregory J., and Robert A. Bloodgood. “Targeting proteins to the ciliary membrane.” Current topics in developmental biology 85 (2008): 115–149.

Pazour, Gregory J., et al. “Polycystin-2 localizes to kidney cilia and the ciliary level is elevated in orpk mice with polycystic kidney disease.” Current Biology 12.11 (2002): R378–R380.

Pazour, Gregory J., et al. “Proteomic analysis of a eukaryotic cilium.” J Cell Biol 170.1 (2005): 103–113.

Pazour, Gregory J., and Joel L. Rosenbaum. “Intraflagellar transport and cilia-dependent diseases.” Trends in cell biology 12.12 (2002): 551–555.

Phua, Siew Cheng, et al. “Dynamic remodeling of membrane composition drives cell cycle through primary cilia excision.” Cell 168.1-2 (2017): 264-279.

Praetorius, Helle A. “The primary cilium as sensor of fluid flow: new building blocks to the model. A review in the theme: cell signaling: proteins, pathways and mechanisms.” American Journal of Physiology-Cell Physiology 308.3 (2014): C198–C208.

Praetorius, Helle A., and Kenneth R. Spring. “The renal cell primary cilium functions as a flow sensor.” Current opinion in nephrology and hypertension 12.5 (2003): 517–520.

Poole, C. Anthony, Michael H. Flint, and Brent W. Beaumont. “Analysis of the morphology and function of primary cilia in connective tissues: a cellular cybernetic probe?.” Cell motility 5.3 (1985): 175–193.

Quan, Tingwei, Shaoqun Zeng, and Zhenli Huang. “Localization capability and limitation of electron-multiplying charge-coupled, scientific complementary metal-oxide semiconductor, and charge-coupled devices for superresolution imaging.” Journal of biomedical optics 15.6 (2010): 066005.

Reese, T. S. “Olfactory cilia in the frog.” The Journal of cell biology 25.2 (1965): 209-230.

Robbins, Mark Stanford, and Benjamin James Hadwen. “The noise performance of electron multiplying charge-coupled devices.” IEEE transactions on Electron Devices 50.5 (2003): 1227–1232.

Rogowski, Michaela, Dirk Scholz, and Stefan Geimer. “Electron microscopy of flagella, primary cilia, and intraflagellar transport in flat-embedded cells.” Methods in enzymology. Vol. 524. Academic Press, 2013. 243–263.

Rohatgi, Rajat, Ljiljana Milenkovic, and Matthew P. Scott. “Patched1 regulates hedgehog signaling at the primary cilium.” Science 317.5836 (2007): 372–376.

Rosenbaum, Joel L., and George B. Witman. “Intraflagellar transport.” Nature reviews Molecular cell biology 3.11 (2002): 813.

Rosenzweig, Derek H., et al. “Subunit dissociation and diffusion determine the subcellular localization of rod and cone transducins.” Journal of Neuroscience 27.20 (2007): 5484–5494.

Ross, Alison J., et al. “Disruption of Bardet-Biedl syndrome ciliary proteins perturbs planar cell polarity in vertebrates.” Nature genetics 37.10 (2005): 1135.

Ruba, Andrew, et al. “Obtaining 3D Super-resolution Information from 2D Super-resolution Images through a 2D-to-3D Transformation Algorithm.” bioRxiv (2017): 188060.

Ruba, Andrew, et al. “Reply to ‘Deconstructing transport-distribution reconstruction in the nuclear-pore complex’.” Nature structural & molecular biology 25.12 (2018): 1062.

Ruba, Andrew, Wangxi Luo, and Weidong Yang. “Application of High-speed Super-resolution SPEED Microscopy in Live Primary Cilium.” Journal of visualized experiments: JoVE 131 (2018).

Scholey, Jonathan M., and Kathryn V. Anderson. “Intraflagellar transport and cilium-based signaling.” Cell 125.3 (2006): 439–442.

Shida, Toshinobu, et al. “The major α-tubulin K40 acetyltransferase αTAT1 promotes rapid ciliogenesis and efficient mechanosensation.” Proceedings of the National Academy of Sciences 107.50 (2010): 21517–21522.

Singh, Jaskirat, Xiaohui Wen, and Suzie J. Scales. “The orphan G protein-coupled receptor Gpr175 (Tpra40) enhances Hedgehog signaling by modulating cAMP levels.” Journal of Biological Chemistry 290.49 (2015): 29663–29675.

Singla, Veena, and Jeremy F. Reiter. “The primary cilium as the cell’s antenna: signaling at a sensory organelle.” science 313.5787 (2006): 629–633.

Sun, Shufeng, et al. “Three-dimensional architecture of epithelial primary cilia.” Proceedings of the National Academy of Sciences 116.19 (2019): 9370–9379.

Vieira, Otilia V., et al. “FAPP2, cilium formation, and compartmentalization of the apical membrane in polarized Madin–Darby canine kidney (MDCK) cells.” Proceedings of the National Academy of Sciences 103.49 (2006): 18556–18561.

von Kleist, Lisa, et al. “Role of the clathrin terminal domain in regulating coated pit dynamics revealed by small molecule inhibition.” Cell 146.3 (2011): 471–484.

Wang, Jing, et al. “The Arf GAP ASAP1 provides a platform to regulate Arf4- and Rab11–Rab8-mediated ciliary receptor targeting.” The EMBO Journal 31.20 (2012): 4057–4071.

Wang, Juan, et al. “C. elegans ciliated sensory neurons release extracellular vesicles that function in animal communication.” Current Biology 24.5 (2014): 519–525.

Wang, Lei, and Brian D. Dynlacht. “The regulation of cilium assembly and disassembly in development and disease.” Development 145.18 (2018): dev151407.

Wheway, Gabrielle, Liliya Nazlamova, and John T. Hancock. “Signaling through the primary cilium.” Frontiers in Cell and Developmental Biology 6 (2018): 8.

Williams, Corey L., et al. “MKS and NPHP modules cooperate to establish basal body/transition zone membrane associations and ciliary gate function during ciliogenesis.” The Journal of cell biology 192.6 (2011): 1023–1041.

Wingfield, Jenna L., et al. “IFT trains in different stages of assembly queue at the ciliary base for consecutive release into the cilium.” Elife 6 (2017): e26609.

Wloga, Dorota, et al. “Posttranslational modifications of tubulin and cilia.” Cold Spring Harbor perspectives in biology 9.6 (2017): a028159.

Wood, Christopher R., and Joel L. Rosenbaum. “Proteins of the ciliary axoneme are found on cytoplasmic membrane vesicles during growth of cilia.” Current Biology 24.10 (2014): 1114–1120.

Yang, T. Tony, et al. “Superresolution pattern recognition reveals the architectural map of the ciliary transition zone.” Scientific reports 5 (2015): 14096.

Ye, Fan, Andrew R. Nager, and Maxence V. Nachury. “BBSome trains remove activated GPCRs from cilia by enabling passage through the transition zone.” J Cell Biol 217.5 (2018): 1847-1868.

Ye, Fan, et al. “Single molecule imaging reveals a major role for diffusion in the exploration of ciliary space by signaling receptors.” Elife 2 (2013): e00654.

Yoder, Bradley K., Xiaoying Hou, and Lisa M. Guay-Woodford. “The polycystic kidney disease proteins, polycystin-1, polycystin-2, polaris, and cystin, are co-localized in renal cilia.” Journal of the American Society of Nephrology 13.10 (2002): 2508–2516.

Yoshimura, Shin-ichiro, et al. “Functional dissection of Rab GTPases involved in primary cilium formation.” The Journal of cell biology 178.3 (2007): 363–369.

