## Supplemental Figures for "The Ciliary Lumen Accommodates Passive Diffusion and Vesicle Trafficking in Cytoplasmic-Ciliary Transport"

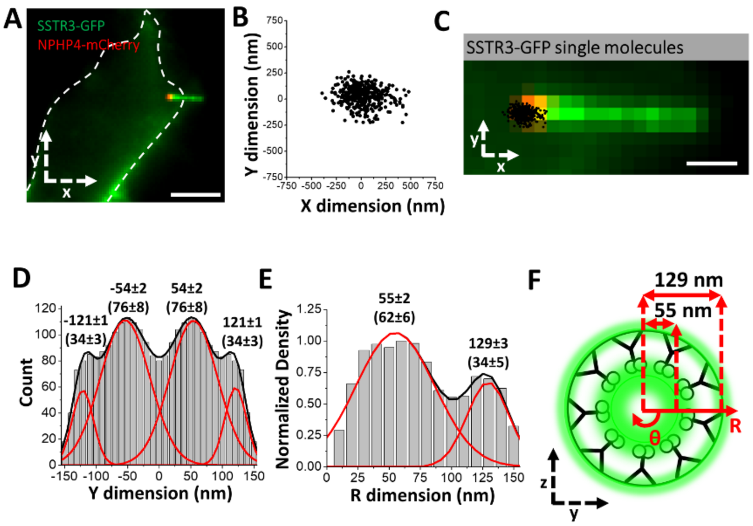


**Supplemental Figure 1.** ***Data transformation process for obtaining 3D density histograms.*** **A)** Epifluorescence microscopy image of live cells co-expressing SSTR3-GFP (green) and NPHP4-mCherry (red) (scale bar: 5 μm). The dashed white line represents the cell border. **B)** Single-molecule SSTR3-GFP locations from the primary cilium in (A). **C)** Enlarged image of the primary cilium overlaid with single molecule SSTR3-GFP locations (black dots). Scale bar: 1 μm. Single-molecule data and epi-fluorescence images were rotated together clockwise, so that TZ is parallel to the x dimension to maintain consistency with subsequent data analyses. **D)** Y-dimensional histogram of the 2D single-molecule SSTR3-GFP locations shown in (B) and fitted by Gaussian functions, generating peak positions and Gaussian widths (in parentheses). Histogram was left-right average due to the bi-lateral symmetry of primary cilia. **E)** Conversion of the 2D data in (D) to a 3D density distribution along the R dimension in the cylindrical system for SSTR3-GFP in the TZ. **F)** Graphical representation of the histogram in (E) and cross-sectional view of SSTR3's two distinct transport routes (green clouds) in the TZ overlaid with the ultrastructure of TZ (grey).

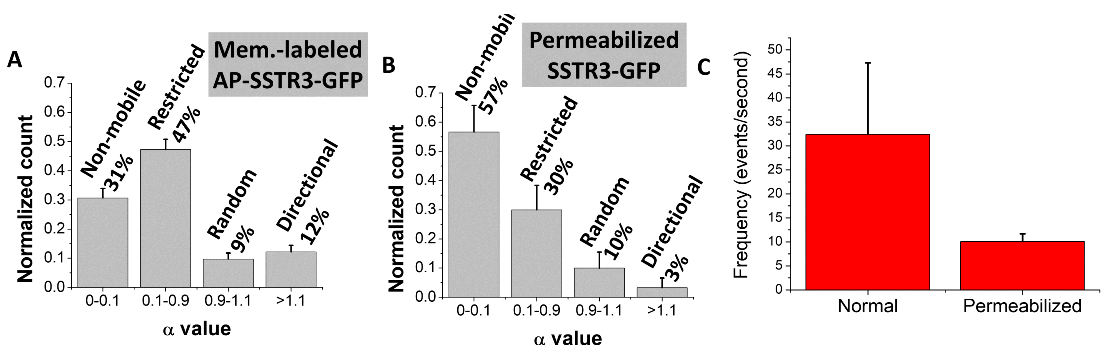


**Supplemental Figure 2.** ***Mean squared displacement analysis for membrane labeled AP-SSTR3-GFP in normal cells and SSTR3-GFP in permeabilized cells as well as single molecule frequency for SSTR3-GFP in normal and permeabilized cells.*** A) Histogram of α values for membrane-labeled AP-SSTR3-GFP trajectories (12% directional movement, n = 205). B) Histogram of α values for SSTR3-GFP trajectories in permeabilized cells (3% directional movement, n = 30). C) Frequency of SSTR3 molecules in the transition zone of live (cilia n = 6) and permeabilized cells (cilia n = 5).

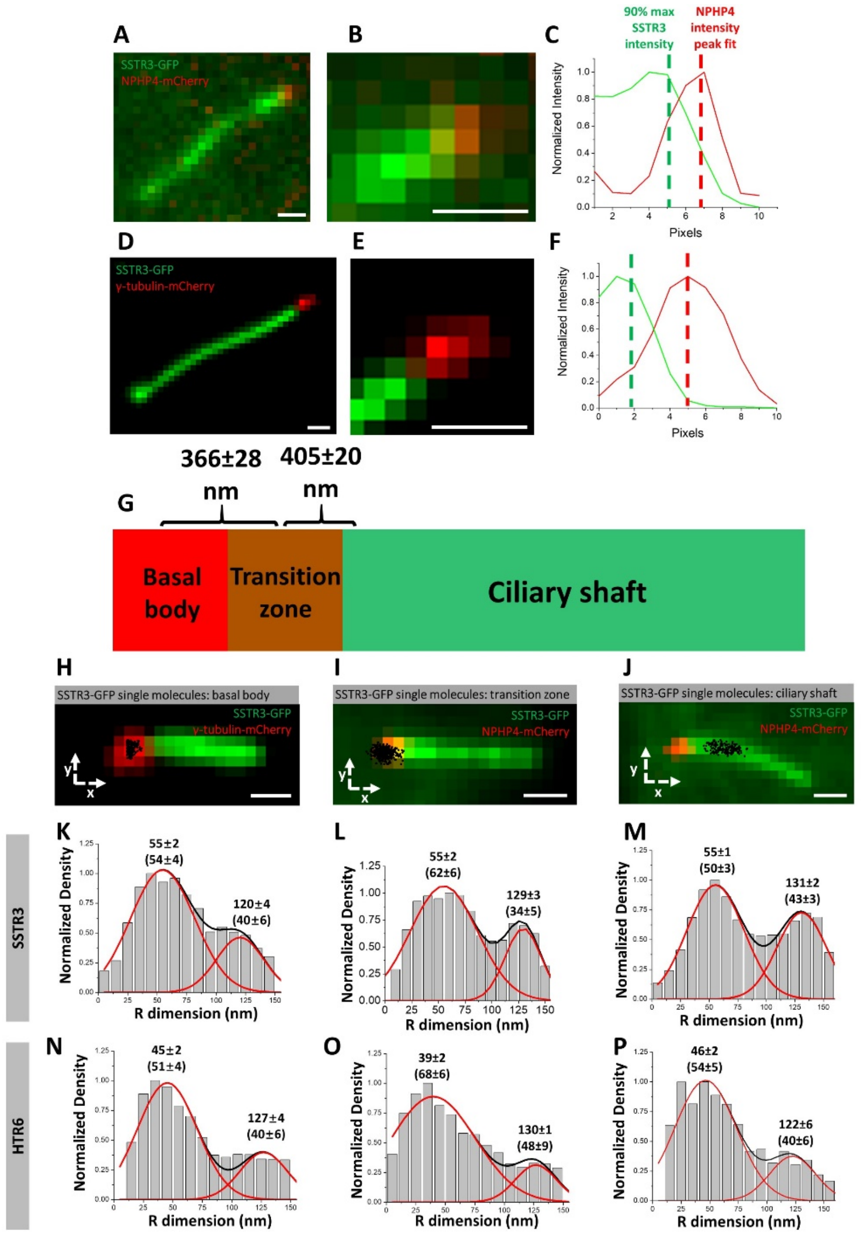


**Supplemental Figure 3.** ***The outer and inner transport routes for HTR6 parallels SSTR3 and is continuous in the basal body, transition zone, and ciliary shaft.*** **A)** Epifluorescence image of SSTR3-GFP (green) and NPHP4-mCherry (red). Scale bar: 1 µm. **B)** Enlarged version of (A). Scale bar: 1 µm. **C)** The intensity plot showing the distance between the peak of NPHP4-mCherry and 90% of the maximum SSTR3-GFP intensity. **D)** Epifluorescence image of SSTR3-GFP (green) and γ-tubulin-mCherry (red).

Scale bar: 1 μm. **E)** Enlarged version of (D). Scale bar: 1 μm. **F)** The intensity plot showing the distance between the peak of γ-tubulin-mCherry and 90% of the maximum SSTR3-GFP intensity. **G)** Schematic of the organization of the basal body, transition zone, and ciliary shaft. **H)** Enlarged image of the primary cilium marked with SSTR3-GFP (green) and γ-tubulin-mCherry (red) and overlaid with single-molecule SSTR3-GFP locations (black dots). Scale bar: 1 μm. **I)** Same as (H) except NPHP4-mCherry (red). **J)** Same as (I). **K), L), and M)** 3D transformed density histogram for SSTR3 at the basal body, transition zone, and ciliary shaft, respectively. **N), O), and P)** Same as (K), (L), and (M) except with HTR6.

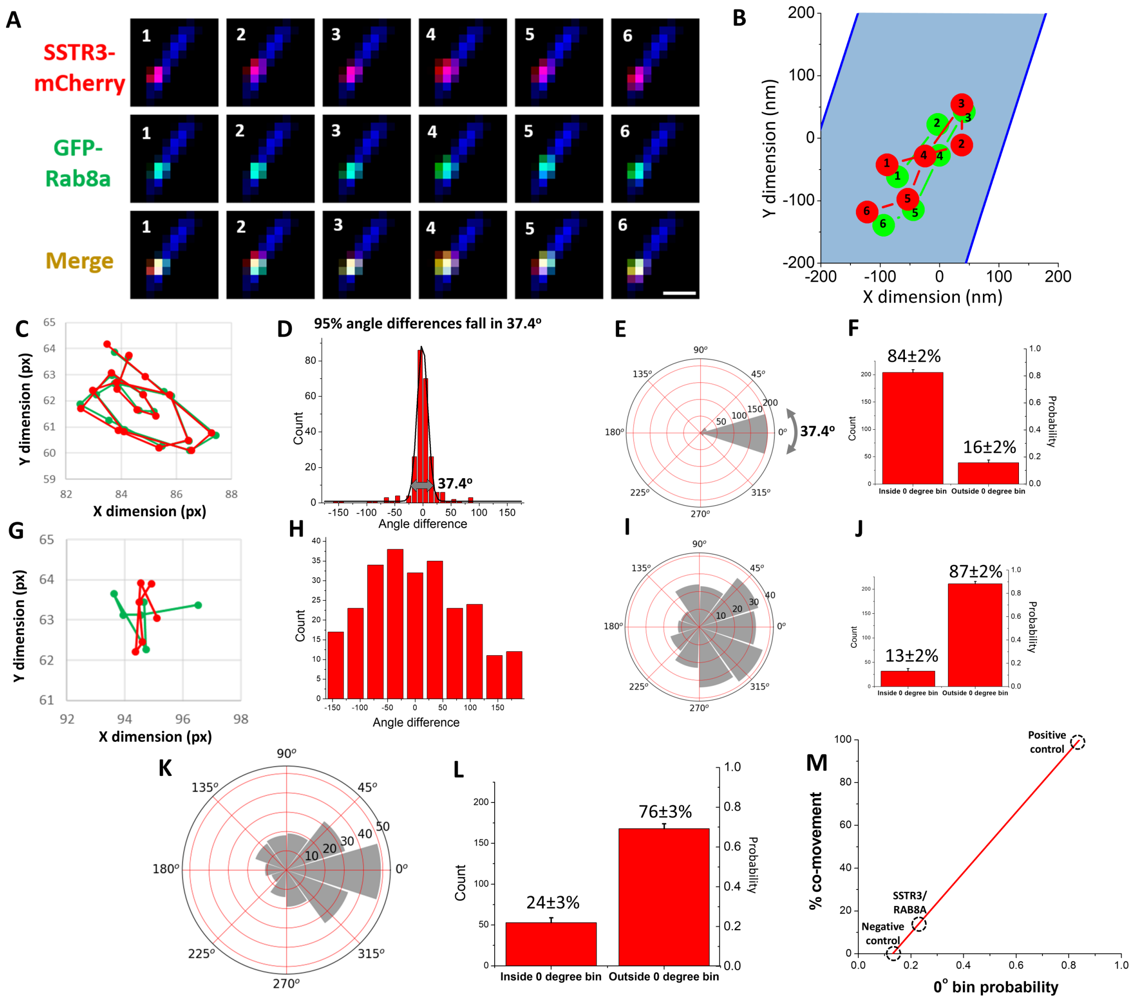


**Supplemental Figure 4.** ***RAB8A shows co-movement with SSTR3 at the ciliary base.***

**A)** Montage of a typical SSTR3-mCherry and GFP-RAB8A single-molecule co-moving event appearing simultaneously and respectively in red and green detection channels near the base of the primary cilium. Scale bar: 1 µm. **B)** Plot of the single-molecule trajectories of SSTR3-mCherry (red) and GFP-RAB8A (green) from (A). Numbers denote frame number. **C)** Single molecule fitting and channel alignment of trajectory in Video 2. **D)** Histogram and peak fitting of all the angle differences in the positive control video. **E)** Radial histogram with the $0^{o}$ bin sized to encompass 95% of the positive control angle differences. **F)** Bar graph showing ratio of angle differences that fall in and out of the 0$^{o}$ bin. **G)-J)** Same as (C)-(F) except with co-tracking of negative control single molecule trajectories. **K)** Radial histogram of the angle difference between each step of all co-appearing SSTR3-mCherry and GFP-RAB8A trajectories that passed selection criteria (Methods). **L)** Bar graph showing the proportion of trajectories that fell in and out of the co-moving bin as defined by the positive control. **M)** Standard curve was calculated by sampling different ratios of trajectories from positive and negative controls which was repeated 1,000 times.

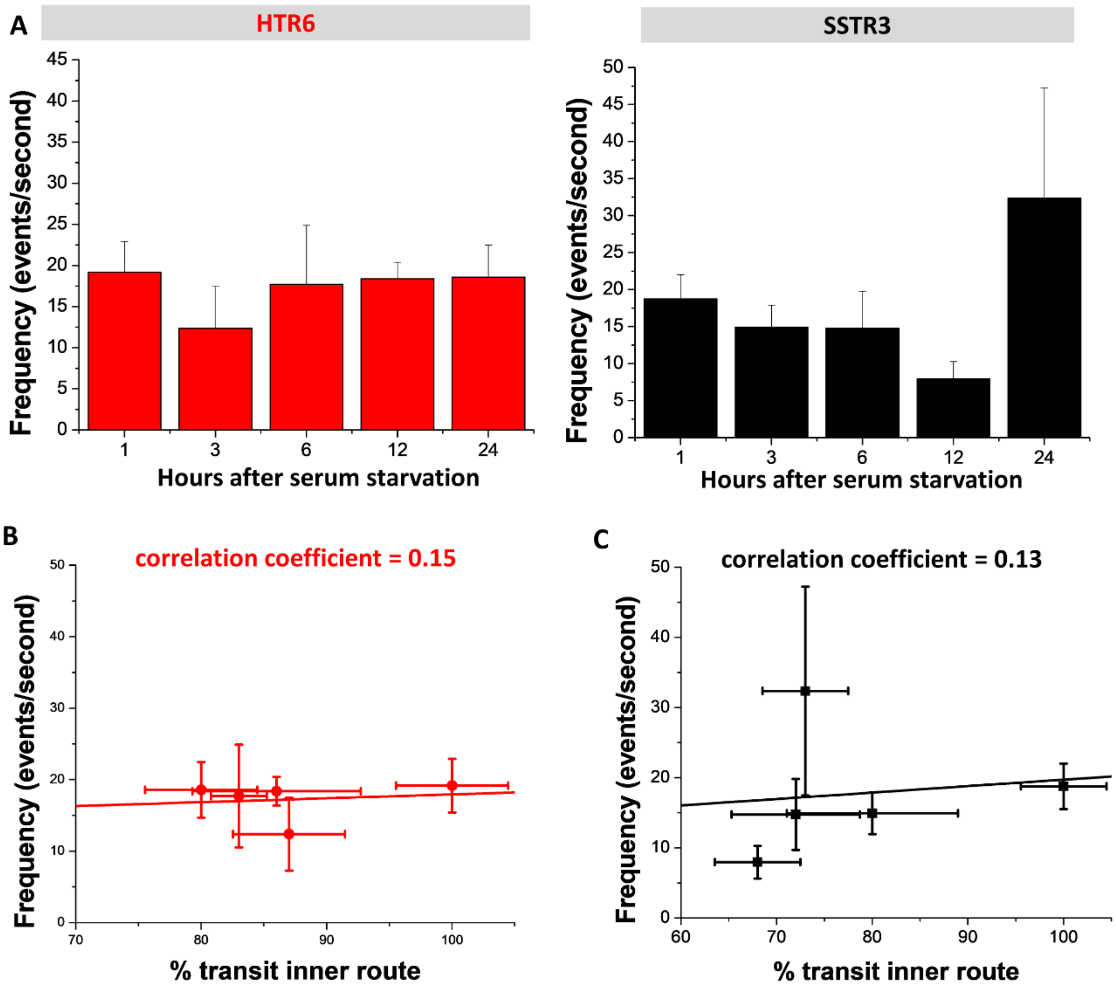


**Supplemental Figure 5.** ***Single molecule frequency does not correlate with transport in the inner route.*** **A)** Single molecule frequency for both SSTR3 and HTR6 at different times after serum starvation. **B)** Single molecule frequency vs. percent transport in inner route for HTR6 with Pearson's correlation coefficient. **C)** Same as (B) with SSTR3.

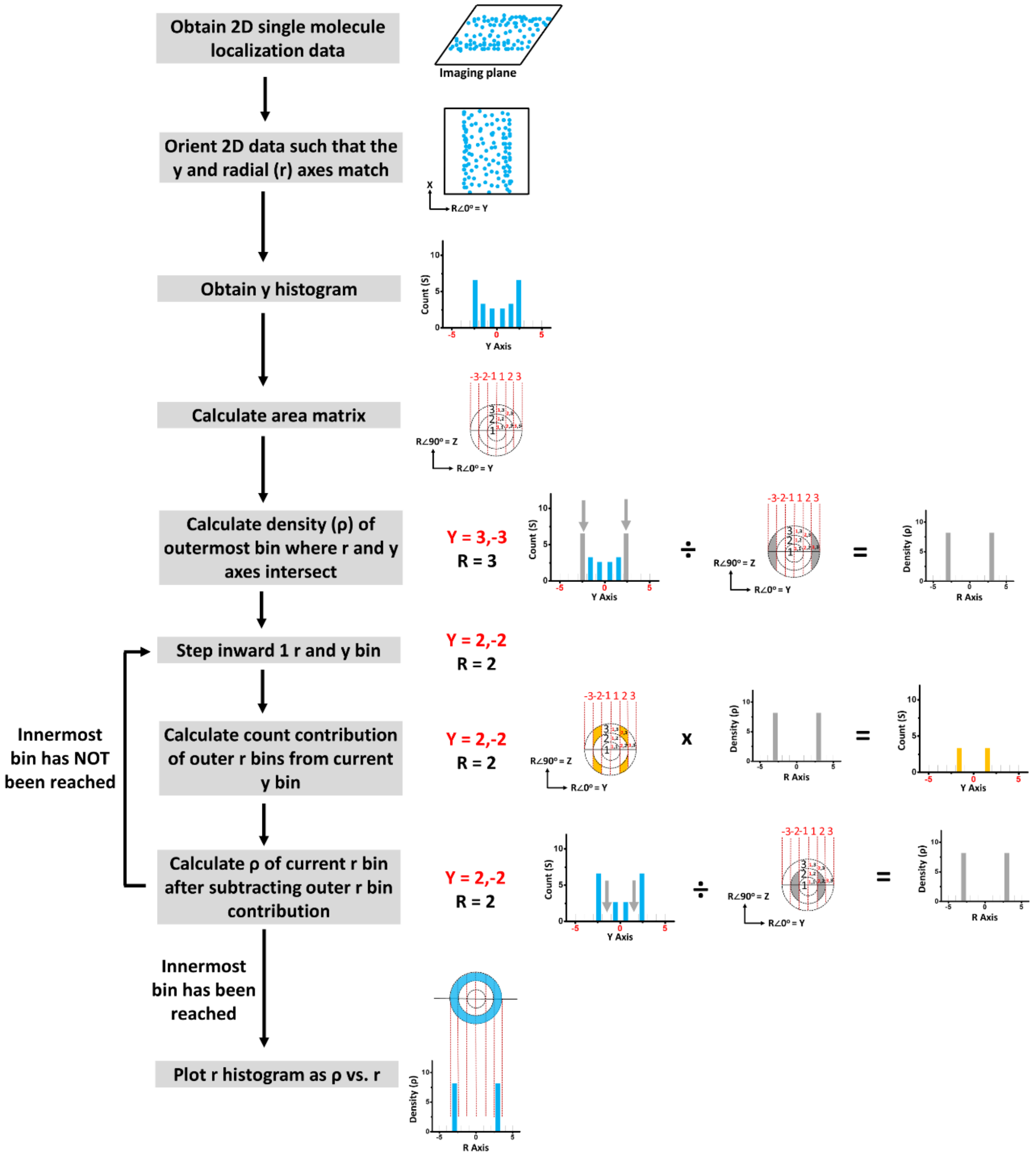


**Supplemental Figure 6.** *Flowchart outlining the conversion of 2D single molecule locations to 3D density histogram.*

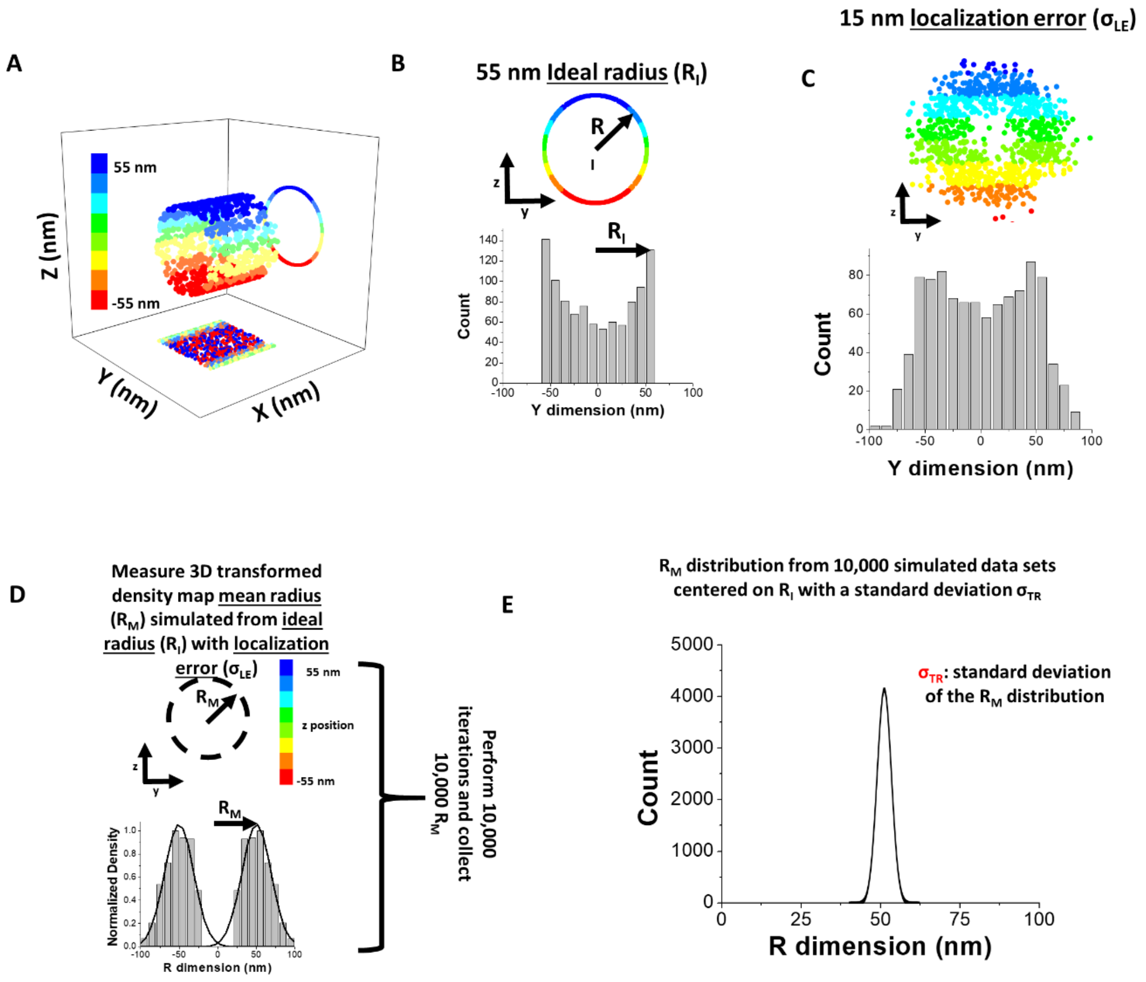


**Supplemental Figure 7.** ***Demonstration of the simulation process.*** **A)** 55 nm ring structure simulated in three dimensions to mimic the SSTR3 inner route. **B)** Y dimensional histogram of (A). **C)** Same as (B) with experimentally derived 15 nm single molecule localization precision. **D)** Calculation of mean peak location following 2D-to-3D transformation of (C). The process is repeated 10,000 times. **E)** Histogram of the mean peak locations derived from (A)-(D). The standard deviation of this distribution corresponds to the spread of possible peak locations given the amount of single molecule locations and single molecule localization precision. We define this standard deviation as $\sigma_{TR}$ or error in determining the mean peak location from the 3D density histogram.

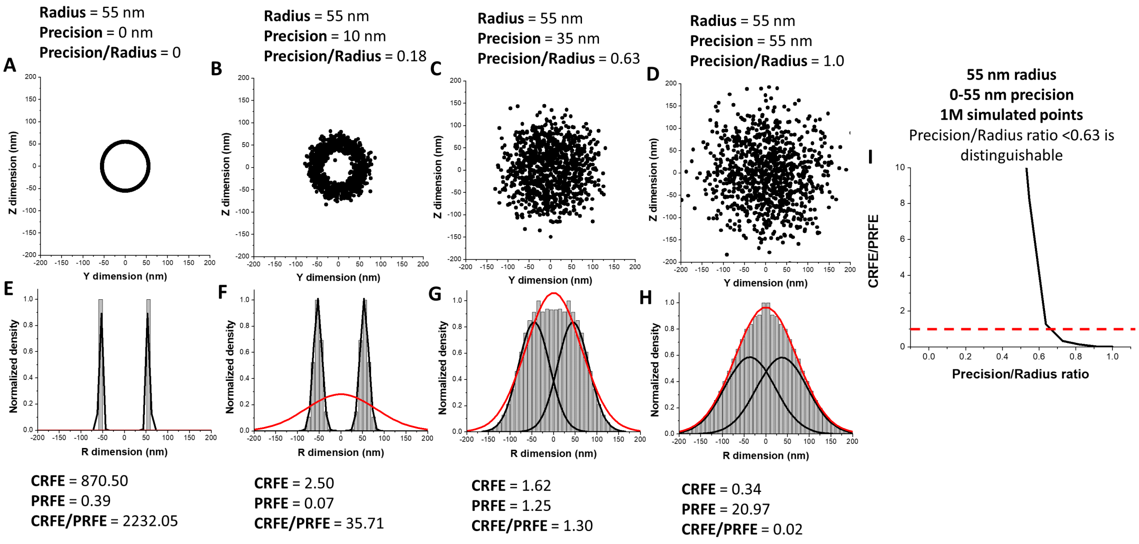


**Supplemental Figure 8.** ***Single molecule localization precision affects the ability to distinguish transport routes in the 3D density histogram.*** **A)-D)** Y,Z simulated locations with varying amounts of simulated single molecule localization precision. **E)-H)** 3D histograms corresponding to the Y,Z locations in (A)-(D) respectively. Central route fitting error (CRFE) compared to peripheral route fitting error (PRFE) is used to determine the ability to distinguish transport routes given varying levels of simulated localization precision. **I)** CRFE/PRFE ratio vs. precision/radius ratio showing a threshold of 0.63, beyond which a peripheral route is no longer distinguishable.

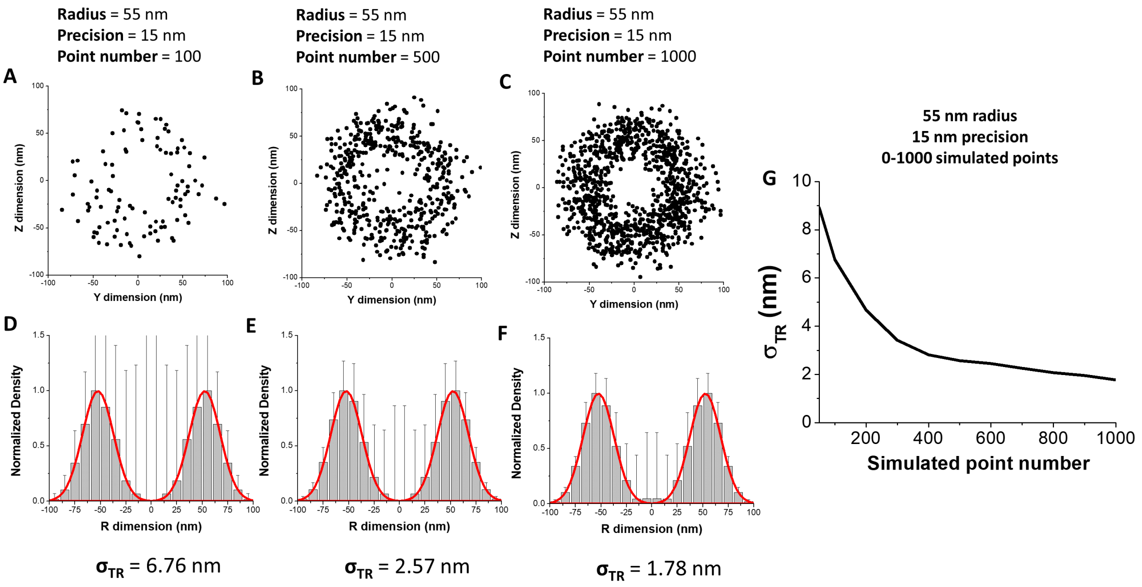


**Supplemental Figure 9.** ***The number of single molecule localizations affects the ability to distinguish transport routes in the 3D density histogram.*** **A)-C)** Y,Z simulated locations with varying amounts of simulated single molecule localizations. **D)-F)** 3D histograms corresponding to (A)-(C). Error bars represent the standard error of each bar in the histogram. **G)** 3D transport route localization precision ($\sigma_{TR}$) vs. the number of simulated localizations.

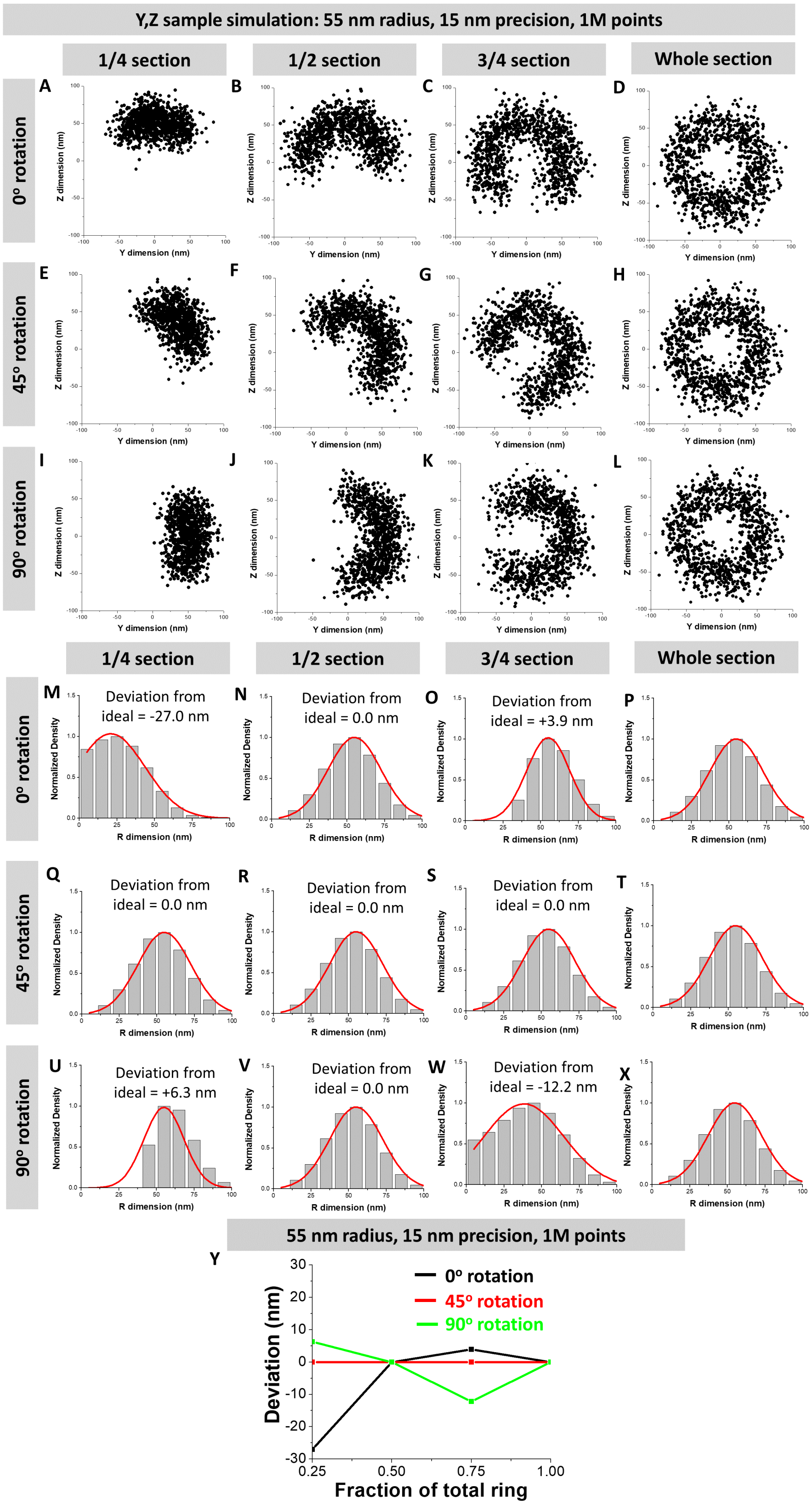
Y,Z sample simulation: 55 nm radius, 15 nm precision, 1M points
1/4 section
1/2 section
3/4 section
Whole section
0º rotation
45º rotation
90º rotation
Y dimension (nm)
Z dimension (nm)
Normalized Density
R dimension (nm)
Deviation from ideal = -27.0 nm
Deviation from ideal = 0.0 nm
Deviation from ideal = +3.9 nm
Deviation from ideal = 0.0 nm
Deviation from ideal = 0.0 nm
Deviation from ideal = 0.0 nm
Deviation from ideal = +6.3 nm
Deviation from ideal = 0.0 nm
Deviation from ideal = -12.2 nm
55 nm radius, 15 nm precision, 1M points
Deviation (nm)
Fraction of total ring
0º rotation
45º rotation
90º rotation

**Supplemental Figure 10.** ***The labeling ratio affects the ability to distinguish transport routes in the 3D density histogram.*** **A)-L)** Y,Z simulated locations for varying ranges of labeling ratio and angle rotation. **M)-X)** 3D transformed histograms from (A)-(L). Depending on the conditions, various degrees of deviation from the actual mean peak location is obtained. **Y)** Deviation from the actual mean peak location vs. labeling ratio over four different degrees of angle rotation.

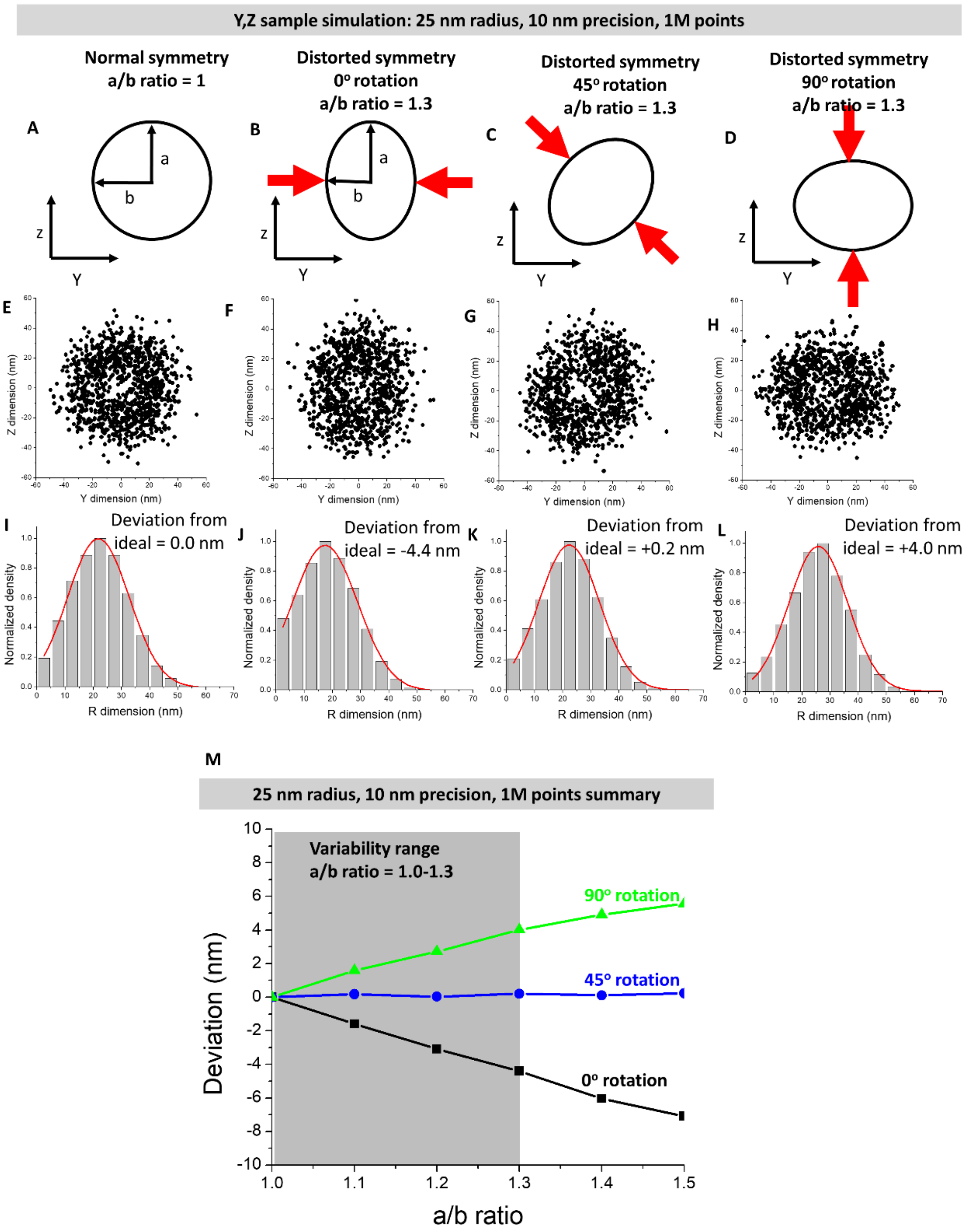


**Supplemental Figure 11.** ***Structural distortion affects the ability to distinguish transport routes in the 3D density histogram.*** **A)-D)** No structural distortion (A) and distortion at various angles (B)-(D) were examined to determine the level of deviation from the expected mean value. **E)-H)** Y,Z simulated data with levels of distortion and rotation corresponding to (A)-(D). **I)-L)** 3D transformed histograms from (E)-(H). **M)** Deviation from the actual peak mean vs. level of distortion at different angles of rotation.

A

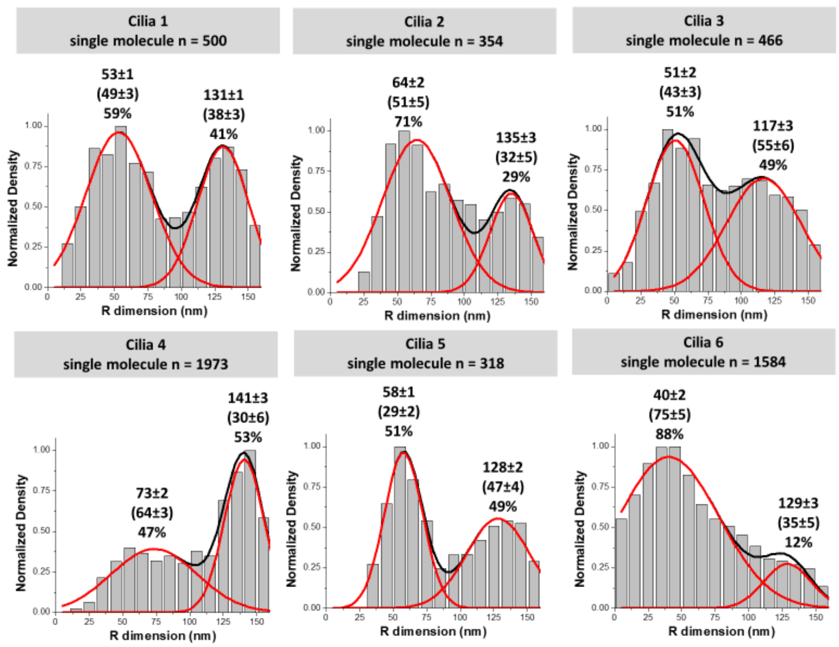


B

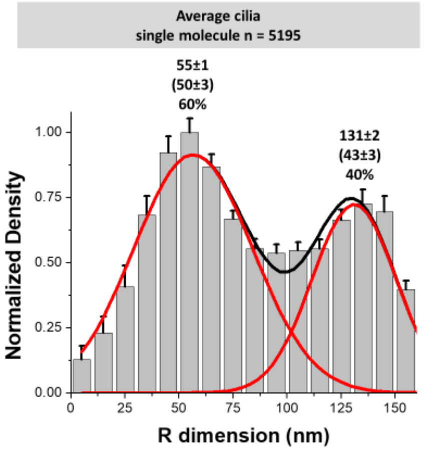


C

cilia n = 6

| Parameter | Standard deviation | Standard error |
|---|---|---|
| Inner route mean (nm) | 11.4 | 4.7 |
| Inner route width (nm) | 16.1 | 6.6 |
| Inner route area (%) | 15.6 | 6.4 |
| Outer route mean (nm) | 8.0 | 3.2 |
| Outer route width (nm) | 9.6 | 3.9 |
| Outer route area (%) | 15.6 | 6.4 |

**Supplemental Figure 12.** ***Variation between different 3D density histograms for SSTR3 in ciliary shaft.*** **A)** 3D transformed histogram from 6 different primary cilia. **B)** Averaged 3D histogram from different primary cilia in (A). **C)** Summary of statistics from (B).

### Materials and Methods

*Tissue culture and transfection*

NIH-3T3 cells or NIH-3T3 cells stably expressing NPHP4-mCherry/IFT20-GFP were grown in DMEM, high glucose, GlutaMAX Supplement (Life Technologies), 10% fetal bovine serum (Fisher Scientific), and 1% penicillin-streptomycin (Thermo Fisher) and split every 2 days to 40-50% confluency. 24 hours prior to imaging, the cells were transferred to glass bottom dishes (MatTek) and grown in OPTIMEM (Life Technologies) to induce growth of primary cilia. Transfection was performed concurrently with induction of primary cilia growth using Transit-LT1 (Mirrus) according to the manufacturer's protocol. Prior to imaging, media was replaced with transport buffer (20 mM HEPES, 110 mM KOAc, 5 mM NaOAc, 2 mM MgOAc, and 1 mM EGTA, pH 7.3). For permeabilization, cells were permeabilized in glass-bottom dishes for 2 min with 30 μg/mL digitonin in transport buffer and washed again with transport buffer. Transport buffer was supplemented with 1.5% polyvinylpyrrolidone. For external membrane labeling, biotin ligase (BirA) and AP-SSTR3-GFP were co-transfected using Transit-LT1 according to the manufacturer's protocol except for the media, which was supplemented with 1 μM biotin. Prior to imaging, cells were incubated with 1 μM AlexaFluor647-streptavidin (Life Technologies) for 30 minutes to cause efficient binding to transiently expressed, biotinylated AP-SSTR3-GFP on the cell surface. Cells were then washed five times with PBS to efficiently wash away unbound AlexaFluor647-streptavidin before placing cells in transport buffer. For Golgicide A inhibition, cells were serum-starved as described above and the media was supplemented with the given concentration of Golgicide A. For somatostatin stimulation, cells were incubated with 10 μM somatostatin for 1 hour prior to imaging. For Pitstop 2 inhibition of somatostatin signaling, cells were incubated with the given concentration of Pitstop 2 for 1 hour prior to addition of somatostatin. For the growth status experiments, cells were serum-starved as described above and imaging was performed at the given time interval post-serum-starvation.

*Plasmids and stable cell lines*

The SSTR3-GFP plasmid was a gift from Kirk Mykytyn (The Ohio State University, College of Medicine). The GFP-IFT43 plasmid was a gift from Kazuhisa Nakayama (Kyoto University, Department of Physiological Chemistry). The IFT20-GFP plasmid was a gift from Gregory Pazour (University of Massachusetts Medical School). Arl13b-mCherry, KIF3A-GFP, SSTR3-mCherry, the IFT20-GFP stable cell line, and the NPHP4-mCherry stable cell line was a gift from Kristen Verhey (University of Michigan). AP-SSTR3-GFP was a gift from Maxence Nachury (Addgene plasmid # 49098), GFP-RAB8A was a gift from Maxence Nachury (Addgene plasmid # 24898), BirA was a gift from Alice Ting (Addgene plasmid #20856), α-tubulin-GFP was a gift from Patricia Wadsworth (Addgene plasmid #12298), and mCherry-Gamma-Tubulin-17 was a gift from Michael Davidson (Addgene plasmid # 55050). The *Chlamydomonas reinhardtii* β-tubulin-GFP and mNG-IFT54 cell lines were a gift from Karl F. Lechtreck (University of Georgia). The *Chlamydomonas reinhardtii* PKD2-GFP cell line was a gift from Kaiyao Huang (Chinese Academy of Sciences). The *Chlamydomonas reinhardtii* KAP-GFP cell line was a gift from Mary Porter (University of Minnesota) through the Chlamydomonas Resource Center (CC-4296).

*Optical setup of SPEED microscopy*

The SPEED microscopy setup includes an Olympus IX81 equipped with a 1.4-NA 100× oil-immersion apochromatic objective (UPLSAPO 100×, Olympus), a 35 mW 633 nm He-Ne laser (Melles Griot), 50 mW solid state 488-nm and 561-nm lasers (Coherent), an on-chip multiplication gain charge-coupled-device camera (Cascade 128+, Roper Scientific) and the Slidebook software package (Intelligent Imaging Innovations) for data acquisition and processing. For individual channel imaging, GFP, mCherry, and Alexa Fluor 647 were excited by 488 nm, 561 nm, and 633 nm lasers, respectively. The fluorescence emissions were collected by the same objective, filtered by a dichroic filter (Di01- R405/488/561/635-25x36, Semrock) and an emission filter (NF01-405/488/561/635-25x5.0, Semrock) and imaged with the above CCD camera operating at either 500 Hz when the 2D-to-3D transformation was performed and 100 Hz when SSTR3/RAB8A cotracking and SSTR3-GFP directionality tracking under somatostatin stimulation.

*Spatial localization of primary cilia and the transition zone*

To mark the transition zone, the centroid of the fluorescent spot for NPHP4-mCherry was used since NPHP4 localizes on the arms of the Y-shaped linkers, which have 9-fold symmetry and are organized in multiple layers. In situations where the tested fluorescently-labeled proteins did not strongly localize to primary cilia, Arl13b-mCherry was used as a ciliary marker. Since Arl13b-mCherry molecules are mobile in primary cilia, they can be quickly pre-photobleached by focusing a 561-nm laser on the ciliary shaft before localization of the NPHP4-mCherry was determined in the transition zone. Since the transition zone may range up to 1000 nm in length along the long axis of primary cilia, only single molecule data collected at ±300 nm from the NPHP4-mCherry centroid was used in the 2D to 3D transformation to ensure only single-molecules moving through the transition zone was collected. To mark the basal body, the above protocol was used except that γ-tubulin-mCherry, the type of tubulin that constitutes the basal body, was used instead of NPHP4-mCherry.

*Single-molecule localization precision*

Single molecule videos were analyzed using Glimpse (Gelles Lab) to determine the XY locations of each single molecule on the imaging plane using 2D Gaussian fitting. The Gaussian width parameter of each single molecule was filtered to ensure only single molecules within the dimensions of the primary cilium were retained for 2D to 3D transformation analysis while the photon count parameter of each single molecule was filtered to ensure adequate precision. The localization precision of immobile fluorescent molecules and moving fluorescent molecules was defined as how precisely the central point of each detected fluorescent diffraction-limited spot was determined. For immobile molecules, the fluorescent spots were fitted to a 2D symmetrical or an elliptical Gaussian function, respectively, and the localization precision was determined by the standard deviation of multiple measurements of the central point. However, for moving molecules, the influence of particle motion during image acquisition should be considered in the determination of localization precision. In detail, the localization precision for moving substrates (σ) was determined by the formula:

$$\sigma = \sqrt{F[\frac{16(s^2 + a^2/12)}{9N} + \frac{8\pi b^2(s^2 + a^2/12)^2}{a^2N^2}]}$$

Where F is equal to 2, N is the photon count, a is the pixel size of the CCD camera, b is the standard deviation of the background in photons per pixel, and s is:

$$s = \sqrt{s_0^2 + \frac{1}{3} D\Delta t}$$

Where $s_0$ is the standard deviation of Gaussian function of the mobile molecules at the focal plane and D is the diffusion coefficient of the mobile molecule (Mortensen et al., 2010, Quan, et al., 2010, Robbins and James, 2003, Deschout et al., 2012). Additionally, $s_0$ and N of single transiting fluorescent molecules were used as selective criteria to ensure that only single molecules with high localization precision within the dimensions of the transition zone were selected. For the single-molecule data used in 2D to 3D transformation, the average single-molecule localization precision for every experiment was summarized in Table 1.

*2D-to-3D density transform algorithm*

The transformation process used to compute the 3D spatial probability density maps of particles transiting through the TZ as has been described in detail in our previous publications and demonstrated again here in Figure S6 (Ma and Yang, 2010, Ruba et al., 2018, Ruba et al., 2017). In short, the 3D spatial locations of molecules transiting through the NPC can be considered in either Cartesian (x, y, z) or cylindrical (x, r, θ) coordinates. In microscopic imaging, the observed 2D spatial distribution of particle localizations is a projection of its actual 3D spatial locations onto the xy plane. Since primary cilia are symmetrical about their long central axis, the 2D spatial distribution is averaged about that axis. The underlying 3D spatial distributions can then be recovered by projection of the measured Cartesian (x, y) coordinates back onto the simplified cylindrical (x, r, constant) coordinates, on the basis of the expected cylindrically symmetrical distribution along the θ direction of the primary cilium.

*Estimation of spatial probability percent for transiting molecules traveling in each transport route*

To calculate the spatial probability of single molecules for a protein with two transport routes, the integrated area of density histogram for each transport route was calculated and expressed as a percentage of the total integrated area of all transport routes to estimate the probability of detecting a single molecule in that given transport route.

*SSTR3 and RAB8A co-movement analysis*

To maximize the chance of detecting of any co-movement between SSTR3 and RAB8A above control levels in live cells, we co-transfected plasmids containing SSTR3-mCherry and GFP-RAB8A into NIH-3T3 cells and used a dual channel filter set to split the individual fluorescence illuminated by simultaneous 561 and 488 nm lasers from each construct onto individual halves of the CCD detector. After alignment of the two halves of the detector, co-moving trajectories in each channel were selected for analysis when at least two consecutive single molecules appeared in each channel, within acceptable WI and localization precision bounds, and no further than 200 nm from each other during each step of the trajectory. This 200 nm requirement was implemented to control for two trajectories were further than a biologically reasonable distance. 200 nm is the approximate diameter of the axoneme plus single molecule localization precision and, therefore, the maximum size of a potential vesicle carrier entering primary cilia. When these requirements were implemented, the positive control – dual labeled 100 nm Tetraspeck beads in 55% glycerol to mimic approximate diffusion constant of *in vivo* molecules - co-movements were highly correlated and negative control – fluorescein and JF561 dye in 92.5% glycerol - co-movements where largely uncorrelated. Glycerol was used to adjust the viscosity of the media and, thus, the diffusion coefficient of the above particles using the following equation: $D = \frac{kT}{6\pi\eta R}$ where D is the diffusion coefficient, k is the Boltzmann constant, T is temperature, η is the viscosity, and R is the radius of the particle. The laser powers for the controls were adjusted to produce localization precisions comparable to experimental conditions. A standard curve was developed from these controls and used to characterize the results *in vivo* (Figure S4 M).

*Calculation of diffusion coefficient and α value*

Calculation of the diffusion coefficient was performed by first plotting each trajectory (>4 frames) on a Mean Square Displacement (MSD) vs. Time (t) plot. The data was fitted with the function $MSD = 4Dt^{\alpha}$ where α is a measurement of graph skewedness. α represents directional movement (or super-diffusion), passive diffusion or sub-diffusion if its value is bigger than 1.1, between 0.9 and 1.1, or smaller than 0.9 respectively.

*Determination of cilia movement, cilia orientation and axial position*

In our experiments, we typically needed two minutes to complete imaging of the entire cilium and collection of ten single-molecule videos (5,000 frames per video and 2 ms per frame) from a primary cilium. Combination of 24-hour serum-starving cell growth and incubation of cells in transport buffer prior to microscopy imaging made the shift of primary cilia in live cells less than 5 nm during the 10 s detection time for each video. As for the cilia orientation, an epi-fluorescence image of Arl13b-mCherry, SSTR3-GFP, HTR6-GFP, or GFP-RAB8A labeled cilium provided a complete image of entire cilium, which clearly indicated the ciliary base, the ciliary tip and the cilia orientation. Then after pre-photobleaching of the tracked protein to locally reduce its concentration, the point-illumination of SPEED microscopy generated 2D super-resolution spatial distribution of protein molecules moving within a range of approximately 1 μm along the ciliary axis in the shaft or TZ of primary cilia. Consequently, the precise location of the middle axis of a primary cilium was obtained by determining the peak position of these 2D super-resolution spatial locations in the ciliary radial dimension with fitting of Gaussian function as well as the Gaussian fitting of the ciliary fluorescence in the image of the ciliary marker.

*Determination of the location of transport route for SSTR3 labeled externally*

The length of the linker was determined by summing the estimated length of each component of the external label: SSTR3 N termius: acceptor peptide:biotin:streptaviding:3xAlexaFluor647 (Howarth and Ting, 2008). The SSTR3 N terminus is 43 amino acids long with a marginal level of disorder according to IUPred. To estimate the length, the average (9.6 nm) between the fully disordered length (43 a.a., 0.4 nm/a.a.) (Erickson, 2009), and fully globular diameter (2 nm) was taken (Ainavarapu et al., 2007, Dosztányi et al., 2005, Erickson, 2009). According to IUPred, the AP domain had a consistent structured prediction. Thus, the globular diameter was used (1.5 nm) (Dosztányi et al., 2005**,** Howarth and Ting, 2008, Erickson, 2009). Using the average length of a C-C bond, biotin's length was estimated to be ~1 nm. Lastly, the globular diameter of tetravalent streptavidin was estimated to be 5 nm (Erickson, 2009). However, three of the four binding domains, on average, are occupied by biotin conjugated AlexaFluor647. Since the average center of fluorescence will be roughly in the middle of the tetravalent streptavidin molecule, 2.5 nm was used for its length. Thus, the total length of the external tag is ~14.6 nm.

*Determination of spatial localization precision of transport route*

As detailed in Figure S7, Monte Carlo simulations were performed where varying numbers of single molecule locations were randomly simulated on an ideal radius ($R_I$). Then, localization error ($\sigma_{LE}$) was added in the y and z dimensions by sampling an error value from a normal distribution with a standard deviation of $\sigma_{LE}$. Subsequently, the 3D transformation algorithm was performed on only the y dimensional data to model the loss of z dimensional information during the 2D microscopy projection process. The peak position of the transformed 3D density histogram was then determined by Gaussian fitting to produce a measured radius ($R_M$) which may deviate from the $R_I$ due to number of simulated locations and simulated localization error. After 10,000 iterations of this process, 10,000 $R_M$ values are collected and the resulting histogram of $R_M$ values can be obtained. After 10,000 iterations, the mean of the $R_M$ values converges on $R_I$, the ideal radius from which all the simulations were originally sampled, while the standard deviation of the $R_M$ values varies based largely on the number of simulated locations and $\sigma_{LE}$. As shown in Table 1, the localization precision and precision/radius ratio of each transport route in primary cilia is high enough to distinguish it and make biologically relevant conclusions.

In addition, all studies indicate that primary cilia have two states of protein transport: 1) ciliary growth state (ciliogenesis) and 2) maintenance state. The process of ciliary growth takes place over the 24 hours of serum starvation. Thus, the 2 minute imaging time of our approach should be sufficient to distinguish even the smallest transition states of protein transport in primary cilia.

*Statistics*

Experimental measurements were reported as mean ± standard error of the mean unless otherwise noted.

*Code availability*

The code for the simulations, 2D-to-3D transformation, and sample and experimental data are available at https://github.com/andrewruba/YangLab.
